# Characterization of *Plasmodium falciparum* NEDD8 and identification of cullins as its substrates

**DOI:** 10.1101/2020.07.21.213488

**Authors:** Manish Bhattacharjee, Navin Adhikari, Renu Sudhakar, Zeba Rizvi, Divya Das, R Palanimurugan, Puran Singh Sijwali

**Author notes:** To whom correspondence should be addressed: Puran Singh Sijwali, CSIR-Centre for Cellular and Molecular Biology, Habsiguda, Uppal Road, Hyderabad-500007, TS, India, (o), /27160311. These authors equally contributed to the study (Manish Bhattacharjee, Navin Adhikari).

## Abstract

A variety of post-translational modifications of *Plasmodium falciparum* proteins, including phosphorylation and ubiquitination, are shown to have key regulatory roles. The neural precursor cell expressed developmentally downregulated protein 8 (NEDD8) is a ubiquitin-like modifier of cullin-RING E3 ubiquitin ligases, which regulate diverse cellular processes, including the cell-cycle. Although neddylation pathway is conserved in eukaryotes, it is yet to be characterized in Plasmodium and related apicomplexan parasites. Towards studying the neddylation pathway in malaria parasites, we characterized *P. falciparum* NEDD8 (PfNEDD8) and identified cullins as its physiological substrates. PfNEDD8 is a 76 amino acid residue protein without the C-terminal tail, indicating that it can be readily conjugated. The wild type and mutant (Gly75Gly76 mutated to Ala75Ala76) PfNEDD8 were expressed in *P. falciparum*. Western blot of wild type PfNEDD8-expressing parasites indicated multiple high molecular weight conjugates, which were absent in the parasites expressing the mutant, indicating conjugation of NEDD8 to proteins through Gly76. Immunoprecipitation followed by mass spectrometry of wild type PfNEDD8-expressing parasites identified several proteins, including two putative cullins. Furthermore, we expressed PfNEDD8 in mutant *S. cerevisiae* strains that lacked endogenous NEDD8 (Δ*rub*1) or NEDD8 conjugating E2 enzyme (Δ*Ubc12*). The western blot of complemented strains and mass spectrometry of PfNEDD8 immunoprecipitate showed conjugation of PfNEDD8 to *S. cerevisiae* cullin cdc53, demonstrating functional conservation and cullins as the physiological substrates of PfNEDD8. The characterization of PfNEDD8 and identification of cullins as its substrates make ground for investigation of specific roles and drug target potential of neddylation pathway in malaria parasites.

## INTRODUCTION

Post-translational modifications (PTM) of proteins such as phosphorylation, methylation, acetylation, glycosylation, ubiquitylation, etc. are key regulators of a plethora of cellular processes, as these modifications cause rapid and dynamic changes to the functions of proteins. Ubiquitin is a major post-translational modifier of proteins in eukaryotes. Mature ubiquitin has 76 amino acid residues arranged into a “beta-grasp” fold, which consists of an α-helix surrounded by a five-strand antiparallel beta sheet [1]. Several protein modifiers are structurally similar to ubiquitin, and are collectively known as ubiquitin-like modifiers (ULMs). Some widely studied ULMs include NEDD8, SUMO, ISG15 and Atg8 [2]. ULMs are major post-translational regulator of protein function by a variety of ways, including protein stability, cellular location, interaction with other proteins, transcriptional regulation, DNA repair and RNA splicing [3,4][5]. ULMs undergo covalent conjugation onto the substrate proteins through a highly regulated cascade of enzymatic reactions, which involves enzymes specific to each ULM.

*Plasmodium falciparum* is the most virulent malaria parasite of humans; the other *Plasmodium* species causing human malaria are *Plasmodium vivax, Plasmodium malariae, Plasmodium ovale* and *Plasmodium knowlesi*. Despite ongoing efforts and advancements to reduce the global burden of malaria, it infected 228 million people, with 405,000 deaths worldwide in 2018 [6]. With increasing resistance of the parasite to most of the commonly used drugs and lack of an effective vaccine, malaria elimination appears to be an elusive goal [7–11]. Hence, there is a need to develop next generation of antimalarials, preferably against novel drug targets. The pathways of ULMs could be promising targets, as each pathway involves multiple enzymes to conjugate the ULM to target proteins, which could be part of multiple cellular processes [12]. Genome wide analysis of the human malaria parasites have revealed the presence of multiple putative ULMs and the associated pathway proteins [13]. The ubiquitin proteasome system (UPS) of malaria parasites has been proposed as an attractive drug target, as inhibitors of the proteasome have been shown to cause parasite death at multiple stages [15,16]. Recent studies demonstrate feasibility of selective inhibition of parasite proteasome, which also had potent effect on artemisinin resistant parasites [17]. *Plasmodium* sumoylation has been shown to be an important regulator of oxidative stress response during erythrocytic stages [18]. *Plasmodium* Atg8 has been shown to be associated with punctate structures and required for apicoplast biogenesis [19–23]. Most of our knowledge on NEDD8 comes from higher eukaryotes and *S. cerevisiae*, whereas the NEDD8 of eukaryotic pathogens like malaria parasites has not been characterized yet.

NEDD8 is synthesized as a precursor, requiring processing by a C-terminal hydrolase, which results in the generation of Gly76 at the C-terminus [24,25]. The conjugation of NEDD8 to substrate proteins is known as neddylation in which Gly76 is conjugated to the ε-amino group of a specific Lys residue in the substrate protein via an iso-peptide bond. Like ubiquitylation, it involves activation of NEDD8 by an activating enzyme E1 called the NEDD8-activating enzyme (NAE), transfer of NEDD8 from E1 to the active site cysteine of the conjugating enzyme E2, and finally conjugation of NEDD8 from the charged E2 to the substrate protein [26–28]. It is not clear if E2 can directly conjugate NEDD8 to the substrate without the involvement of an E3 ligase. However, several ubiquitin E3 ligases have been shown to facilitate self-neddylation or neddylation of other proteins, thereby potentially acting as NEDD8 E3 ligases [29,30]. For example, mdm2 has been shown to not only mediate neddylation of p53 and p73 but also mediates self-neddylation [31]. Human NAE is a heterodimer of ubiquitin-activating enzyme 3 (UBA3) and amyloid precursor protein-binding protein 1 (APPBP1). UBA3 functions as the catalytic subunit and APPBP1 functions as the regulatory subunit. Human and *S. cerevisiae* NEDD8 E2 is called ubiquitin conjugating enzyme 12 (Ubc12). In spite of the high sequence identity between human NEDD8 and ubiquitin (60%), certain unique residues in ubiquitin and NEDD8 ensure their specific recognition and conjugation by their respective enzymatic machineries [32].

Neddylation is an essential process in all the eukaryotes studied so far [33–39] except *S. cerevisiae* [40]. The outcome of neddylation can vary from change in sub-cellular localization [41], interactions [42], activation [43] to conformational changes [44]. The best defined and conserved function of neddylation is activation of Cullin-RING E3 ubiquitin ligases (CRLs), including SCF (Skp1, Cullins, F-box proteins) and APC (Anaphase-promoting complex), which play crucial roles in cell-cycle regulation [44,45], hypoxia signaling [46], DNA damage repair [47]. The neddylation of cullin in CRLs enhances ubiquitin ligase activity [48–50]. Neddylation of histone H4 at DNA damage sites serves as a signal to recruit the ubiquitylation machinery for repair of the damaged DNA [29]. Neddylation of epidermal growth factor receptor (EGFR) has been shown to be an important upstream signal leading to increased ubiquitylation and endocytic internalization [51]. Many forms of malignancies have also been shown to have chronic dependence of neddylation to thrive, as cancer cells are heavily dependent on signals that promote cell-cycle progression, and many of these regulatory proteins are neddylation substrates [25,46]. In case of breast cancer, neddylation of breast cancer-associated protein 3 has been shown to recruit a class III histone deacetylase that represses NFκB-dependent transcription [42]. It has also been demonstrated that higher neddylated forms of key oncogenic hallmarks LKB1 and Akt stabilize them, which in turn induces metabolic disruptions, progressively leading to liver cancer [53]. Hence, NAE has been explored as a target for development of anti-cancer drugs, and its inhibitor MLN4924 has shown promise as an anti-cancer drug [52,55,56].

Bioinformatics analysis of the genomes of protozoan parasites has predicted the presence of NEDD8 and some components of the neddylation pathway [12]. The only experimental report on the neddylation pathway of a protozoan parasite is of *Trypanosoma brucei*, which highlighted a few atypical features of its NEDD8 in comparison to NEDD8 from higher eukaryotes [57]. *Plasmodium* neddylation has been explored from the point of de-neddylation, and PfUCHL3 and PfUCH37 were demonstrated to have de-ubiquitylase as well as de-neddylase activities. Interestingly, these studies also revealed that among its homologs in higher eukaryotes, only PfUCH37 has de-neddylase and de-ubiquitylase activities. However, PfUCHL3, the essential one among the two, was unable to de-neddylate cullin, suggesting the presence of other de-neddylase(s) in *Plasmodium* [58–61]. As neddylation is a very well acknowledged critical regulator of fundamental cellular processes like cell cycle in higher eukaryotes [31,48,50], it is likely to have important roles during parasite development.

Here, we characterize the *Plasmodium falciparum* NEDD8 and demonstrate that *P. falciparum* cullins are NEDD8 substrates. This work lays the foundation for understanding the role of protein neddylation in malaria parasites.

## MATERIALS AND METHODS

*Plasmodium falciparum* D10 strain was obtained from the Malaria Research and Reference Reagent Resource Centre (MR4). MLN4924 was from Adooq Bioscience, and all other biochemicals were from Sigma or Serva unless otherwise mentioned. Plasmid and genomic DNA isolation kits were from MACHEREY-NAGEL, cell culture reagents were from Lonza and Invitrogen, restriction enzymes and DNA modifying enzymes were from New England Biolabs and Thermo scientific. Antibodies were from Cell Signaling Technologies, Santacruz Biotechnology and Thermo Fisher Scientific. PCR reagents were from Thermo Fisher Scientific or New England Biolabs. Ni-NTA resin was from Thermo Fisher Scientific. Human blood was collected from healthy volunteers after written consent under medical supervision at the medical dispensary of the institute according to the protocols approved by the Institutional Ethical Committee of Centre for Cellular and Molecular Biology, India. Yeast extract, peptone, agar and yeast nitrogen base were from HiMedia. Glucose, galactose and amino acids (arginine, histidine, isoleucine, leucine, lysine, methionine, phenylalanine, threonine, tryptophan) were from Sigma. Geneticin (G418) was from Calbiochem. Lithium acetate and PEG were from Sigma and the carrier DNA was from Gibco. *S. cerevisiae* strain BY4741 (*MATa his3Δ1 leu2Δ0 met15Δ0 ura3Δ0*; Source EUROSCARF: FY1769) and plasmid pLE124 were kind gifts from Dr. Palani Murugan Rangasamy.

### Sequence analysis

For identification of *Plasmodium* NEDD8 proteins, human NEDD8 (UniProt ID: Q15843) and *S. cerevisiae* NEDD8 (UniProt ID: Q03919) sequences were used as queries in BLAST searches of the genomes of malaria parasites at PlasmoDB (http://plasmodb.org) and the non-redundant protein sequence database at National Centre for Biotechnology Information (http://www.ncbi.nlm.nih.gov) [62,63]. The BLAST search was performed using BlastP algorithm, with BLOSUM62 as the scoring matrix. The top *P. falciparum* hits were used as queries in reverse BLAST searches against the UniProt database (https://www.uniprot.org/). The hits were analyzed to identify conserved domains and substantiate their authenticity to be NEDD8 proteins using the conserved domain database (http://www.ncbi.nlm.nih.gov/Structure/cdd/wrpsb.cgi). Sequence alignments were performed using the Clustal Omega (https://www.ebi.ac.uk/Tools/msa/clustalo/) or MULTALIN (https://npsa-prabi.ibcp.fr/cgi-bin/npsa_automat.pl?page=/NPSA/npsa_multalin.html) programs, and alignment was edited manually [64]. The sequences of NEDD8 homologs from a diverse range of eukaryotes representing metazoans, plants, fungi and protozoans were obtained from the UniProt database. Since many of the NEDD8 homologs retrieved were c-terminal fusion proteins of ubiquitin or other proteins, only the sequences corresponding to NEDD8 were taken for further analysis. To perform phylogenetic analysis of NEDD8 homologs, we subjected them to structure-based sequence alignment (Expresso) in T-coffee server (http://tcoffee.crg.cat) [65]. The aligned sequences were taken to construct a phylogenetic tree using the maximum likelihood statistical method, with Jones-Taylor-Thornton substitution model in MEGA-X software [66,67].

### Parasite culture

In vitro culture of *P. falciparum* was done according to the protocols approved by the Institutional Biosafety Committee (IBSC) and Institutional Ethics Committee of Centre for Cellular and Molecular Biology. *P. falciparum* D10 strain was cultured in human erythrocytes at 2% haematocrit in the presence of a gas mixture (5% CO_2_, 5% O_2_ and 90% N_2_) in RPMI1640-albumax medium (RPMI1640 with 25 mM HEPES, 2 g/l sodium bicarbonate, 2 g/l glucose, 25 μg/ml gentamicin, 300 mg/l L-glutamine, 0.5 % albumax II) [68]. Synchronous cultures were obtained by treating parasites with 5% D-sorbitol [69]. For isolation of parasites, asynchronous parasite culture or parasite cultures of desired stages were harvested at 5-10% parasitemia by centrifugation at 453g for 5 min at room temperature. The cell pellet was incubated with 5× pellet volume of ice-cold 0.05% saponin (in PBS) for 5 min [19], the lysate was centrifuged at 2465g for 8 min at 4°C, the supernatant was discarded, the pellet was washed twice with ice-cold PBS to remove erythrocyte membranes, and the final parasite pellet was stored at −80°C until further use. For total RNA, the pellet of trophozoite/schizont stage parasites was processed using the Nucleospin^®^ RNA kit (Macherey-Nagel) as instructed by the manufacturer. 5 μg of genomic DNA-free total RNA was used for cDNA synthesis using the Invitrogen Superscript™-III first-strand synthesis system as described by the manufacturer. Genomic DNA was isolated from the pellet of trophozoite/schizont stage parasites using the Nucleospin^®^ tissue kit (Macherey-Nagel) as instructed by the manufacturer.

### Transfection vectors and transfection of *P. falciparum*

The PfNEDD8 (PlasmoDB ID: PF3D7_1313000) coding region was amplified from *P. falciparum* gDNA using Phusion DNA polymerase and specific primers PfHN8-F/PfN8ex-R (Table S1). This fragment would code for HA-tagged PfNEDD8 (HN8). A PfNEDD8 mutant (HN8GGm), with Gly75-Gly76 substituted by Ala-Ala, was amplified from *P. falciparum* gDNA using the primers PfHN8-F/PfN8ggM-R). The PCR fragments were A-tailed, cloned into the pGEMT plasmid, and transformed into DH5α *E. coli* cells to obtain pGEMT-HN8 and pGEMT-HN8GGm plasmids. The plasmids were sequenced to confirm the insert sequence, excised with BglII/XhoI, and subcloned into the similarly digested pNC vector to obtain pNC-HN8 and pNC-HN8GGm plasmids. pNC was derived from PfCENV3 [70], which involved excision of the centromeric region with EcoRI, followed by ligation of the vector backbone. pNC-HN8 and pNC-HN8GGm plasmids were digested with various combinations of restriction enzymes to confirm the presence of different regions. Large quantities of plasmid DNAs for transfection were prepared from the pellets of overnight cultures of pNC-HN8 and pNC-HN8GGm clones using the NucleoBond Xtra Midi kit (Macherey-Nagel) as instructed by the manufacturer.

A freshly thawed culture of *P. falciparum* at ~10% early ring stage parasitemia was used for transfection as has been described earlier [71–73] The required volume of culture (5 ml/transfection) was centrifuged at 453xg for 5 minutes at room temperature, the pellet (~100 μl packed cell volume) was washed with cytomix (120 mM KCl, 0.15 mM CaCl_2_, 2 mM EGTA, 5 mM MgCl_2_, 10 mM K2HPO4/KH2PO4, 25 mM HEPES, pH 7.6) [74,75], mixed with 320 μl of cytomix containing 50-100 μg plasmid DNA (pNC-HN8 or pNC-HN8GGm), the suspension was transferred to a chilled 0.2 cm electroporation cuvette, and pulsed (950 μF, 310 mV, and ∞ resistance) using the Bio Rad GenePulser XCELL. The cuvette content was immediately transferred to a flask with 3 ml culture medium, the flask was gassed and incubated at 37°C for 3 hours. Three hours post-transfection, the culture was centrifuged, supernatant was discarded, and the pellet was resuspended in 5 ml fresh culture medium. On day 2, the culture was expanded with fresh RBCs to adjust parasitemia to ~5%, and selection was started with 1 μg blasticidin/ml. The culture medium was changed every day for 7 days and then after every other day till resistant parasites emerged (2-3 weeks). 25 μl of fresh blood (50% haematocrit) was added every week to the culture until the culture had sufficient resistant parasites to be expanded. Upon emergence of resistant parasites, the cultures were maintained in the presence of 0.5 μg blasticidin/ml, which was withdrawn during experiments.

### Expression and localization of PfNEDD8

To determine expression of wild type PfNEDD8, an asynchronous culture of HN8-expressing parasites was harvested at 10-15% parasitemia as described in the parasite culture section. To determine expression of wild type PfNEDD8 in different stages, a synchronized culture of HN8-expressing parasites was harvested at ring, trophozoite, and schizont stages. To check the expression of mutant PfNEDD8, an asynchronous culture of HN8GGm-expressing parasites was harvested at 10-15% parasitemia. The parasite pellets were resuspended in 3× pellet volume of 10 mM Tris, pH 7.5 and 1× volume of 4×SDS-PAGE sample buffer (1× buffer contains 50 mM Tris-HCl, 20% glycerol, 2% SDS, 1% β-mercaptoethanol, 0.01% (w/v) bromophenol blue, pH 6.8). The resuspension was incubated at 99°C for 10 min, centrifuged at 24000g for 15 minutes, and supernatant was transferred to a fresh tube. Equal amounts of supernatants (~1×10^8^ parasites/lane) were resolved on a 12% SDS-PAGE gel and transferred onto the Immobilon-P membrane. The membrane was incubated with blocking buffer (3% skimmed milk in 10 mM Tris-Cl, 150 mM NaCl, 0.1 % Tween-20, pH-7.4) for one hour at room temperature, followed by with primary antibodies (rabbit anti-HA antibodies at 1/1000 or mouse anti-β-actin antibodies at 1/1000 in blocking buffer) for 1 hour at room temperature. The membranes were washed with TBST (10 mM Tris-Cl, 150 mM NaCl, 0.1%Tween-20, pH-7.4), incubated with appropriate secondary antibodies (HRP-conjugated goat anti-rabbit or HRP-conjugated goat anti-mouse IgG at a dilution of 1/20,000 in blocking buffer) for 1 hour at room temperature. The blots were washed, and signal was developed using the SuperSignal™ West Pico or Femto Chemiluminescent kit (Thermo Fisher Scientific), and recorded on the BioRad ChemiDoc™ MP imaging system.

For localization of wild type PfNEDD8, 200 μl of an asynchronous culture of HN8-expressing parasites was centrifuged at 11000xg for 10 seconds, the pellet was washed with PBS, layered on a poly L-Lysine coated slide for 20 minutes, unbound cells were washed off with PBS, and the immobilized cells were fixed (3% paraformaldehyde and 0.01% glutaraldehyde) for 45 mins. The cells were permeabilized with 0.1% (v/v) Triton X-100 (in PBS) for 30 minutes, blocked with blocking buffer (3% BSA in PBS) for overnight at 4°C, and incubated with rabbit anti-HA antibodies (1/100 in blocking buffer) for 1 hour. The cells were washed with blocking buffer, incubated with secondary antibody (Alexa Fluor 594-conjugated donkey anti-rabbit IgG (at 1/1000 dilution in blocking buffer) for 1 hour, and then incubated with the nuclear stain DAPI for 20 minutes (10 μg/ml in PBS). The slides were air-dried, mounted with ProLong Gold anti-fade reagent, the sample area was covered with a coverslip, and the edges were sealed with nail polish [19]. The slide was observed under 100× objective of the AxioImager.Z1, images were captured with AxioCam, and analysed using the AxioVision LE software.

### Effect of MLN4924 and mutation of PfNEDD8 on neddylation

To assess the effect of a general neddylation inhibitor MLN4924 on *P. falciparum* growth, the wild type *P. falciparum* was cultured with different concentrations of MLN4924. The MLN4924 stock (45.1 mM) was serially diluted 2-fold in 50 μl of culture medium across rows of a 96-well tissue-culture plate. DMSO (0.5%) or 200 nM chloroquine (CQ) was added to the control wells. 50 μl parasite suspension (1-2% ring-infected erythrocytes at 4% hematocrit) was added to each well. The plate was incubated in a modular incubator chamber (Billups-Rothenberg, Inc.) along with the gas mixture at 37°C for 48-50 h. Post-incubation, the culture medium was aspirated and the cells were resuspended in 100 μl of 2% formaldehyde (in PBS) for 48 hours in the dark. 20 μl from each well was transferred to a new plate, mixed with 100 μl of labelling solution (0.1% Triton X-100 in PBS with 10 nM YOYO-1; Thermo Fisher Scientific), and analysed on BD Fortessa FACS Analyzer (λex 491 nm and λem 509 nm). 10,000 cells were counted, and the percent of infected cells was determined. The parasitemia of CQ sample was subtracted from those of DMSO and MLN4924 samples to adjust for the background, and the parasitemias of MLN4924 samples were normalized with that of DMSO sample to calculate % inhibition, which was plotted against MLN4924 concentrations and analysed using nonlinear regression curve fit (GraphPad Prism) to calculate the IC_50_ concentration of MLN4924 for the parasite growth inhibition.

To determine if mutation of the PfNEDD8 C-terminal Gly75Gly76 to Ala-Ala affects neddylation, 20 ml culture aliquots (10% parasitemia) of synchronized HN8-expressing *P. falciparum* parasites and HN8GGm-expressing *P. falciparum* parasites at trophozoite stage (30 hours post-synchronization) were purified and processed for western blotting with anti-HA antibodies as mentioned in the western blotting section.

To determine if MLN4924 affects neddylation, 10 ml aliquots (10% parasitemia) of synchronized HN8-expressing *P. falciparum* parasites at early trophozoite stage (24 hours post-synchronization) were cultured in the presence of 175 μM MLN-4924 or 0.4% DMSO for 4 or 8 hours. At the end of treatment, the parasite morphology was monitored by bright-field microscopy, parasites were purified and processed for western blotting with anti-HA antibodies as mentioned in the western blotting section.

### Production of recombinant PfNEDD8

The PfNEDD8 coding region was amplified from *P. falciparum* cDNA using Phusion DNA polymerase and primers SN8-F/SN8-R. A SUMO-fusion vector (pSMO) was constructed for expression of PfNEDD8 as a C-terminal fusion of SUMO. The *S. cerevisiae* SUMO/SMT-3 (UniProt ID: Q12306) coding region was amplified from the gDNA using Phusion DNA polymerase and primers SMT3-F/SMT3-R. The PCR product was cloned into the NdeI-HindIII site of pET32a vector, replacing the thioredoxin/His/polylinker region. The PfNEDD8 PCR product was cloned into the StuI-HindIII digested pSMO plasmid using in-Fusion^®^ HD Cloning Plus kit (Takara) and transformed into DH5α *E. coli* cells to obtain pSMO-PfNEDD8 plasmid. The clones were sequenced to ensure that there were no errors in the PfNEDD8 gene. The sequence confirmed pSMO-PfNEDD8 plasmid was transformed into Rosetta-origammi *E. coli* cells, which would express PfNEDD8 as a C-terminal fusion of His-SUMO (SN8). A culture of pSMO-PfNEDD8 expression clone was induced with IPTG (1 mM) at OD_600_ of 0.6 for 4 hours at 25°C with shaking at 150 rpm. The culture was harvested and, the pellet was resuspended in lysis buffer (50 mM Tris-Cl, 150 mM NaCl, 10 mM imidazole, pH 8.0; 5 ml/g pellet weight) with lysozyme (1 mg/ml) at 4°C for 30 minutes, the lysate was sonicated for 4 minutes (pulses of 9 seconds ON and OFF at maximum amplitude). The lysate was centrifuged at 20000g for 15 minutes at 4°C, the supernatant was transferred to a new tube and centrifuged again at 35000g for 30 minutes at 4°C. The supernatant was incubated with Ni-NTA resin (preequilibrated with lysis buffer; 0.25 ml resin slurry/1.0 g pellet weight) for 30 min at 4°C. The suspension was transferred to a column, unbound proteins were allowed to pass through, the resin was washed with 50× column volume of the lysis buffer (containing 20-50 mM imidazole), and the protein was eluted with 250 mM imidazole (in lysis buffer). Elution fractions were run on 12% SDS-PAGE, the fractions containing pure protein were pooled and dialysed against 100-fold excess of dialysis buffer (20 mM Tris-Cl, 50 mM NaCl, pH 7.5) using a 3 kDa cut-off dialysis tubing at 4°C for 16 hours, including one change of the buffer after 10 hours. The dialysed sample was concentrated to 1 mg/ml using Ultracel-3k (Millipore) at 4°C, and stored at −30°C till further use.

### In-vitro neddylation assay

Neddylation assay requires NEDD8, NAE, Ubc12, neddylation substrates, ATP and mild reducing conditions [76,77]. Purified recombinant SN8 was used for in-vitro neddylation assay using soluble extract of wild type *P. falciparum* parasites as a source of enzymes and substrates. For preparing parasite extract, a pellet of asynchronous parasites was resuspended in the lysis buffer (20 mM Tris-Cl, pH 7.5) and subjected to 2 cycles of lysis using the Diagenode Bioruptor^®^ for 5 minutes (each cycle involves pulses of 30 seconds ON and 30 seconds OFF). The lysate was centrifuged at 25000g for 30 min at 4°C, the supernatant that contained soluble parasite extract was transferred into a fresh tube, and the protein amount in the extract was estimated using BCA method. The parasite extract was passed twice through fresh Ni-NTA resin to remove any interacting proteins. The in vitro neddylation assay reaction contained 200 μl of reaction buffer (50 mM Tris-Cl, 0.5 mM DTT, MgCl_2_, pH 7.5) containing 100 μg of parasite extract, 1 μg of recombinant SN8 and 5 mM ATP. In parallel, three control reactions were set up: no ATP, parasite extract with ATP but without SN8, SN8 with ATP but without parasite extract. All the reactions were incubated at 37°C for 1 hour with mild shaking. The reactions were either stopped by adding 50 μl of 5× SDS-PAGE sample buffer and used for western blot analysis or directly used for isolation of SN8-conjugates using Ni-NTA resin. Aliquots of reaction samples were processed for western blot analysis using rabbit anti-His antibodies (1:1000 dilution in blocking buffer) and mouse anti-β-actin antibodies (1:1000 dilution in blocking buffer), followed by appropriate secondary antibodies. For isolation of SN8-conjugates, the reaction samples were incubated with Ni-NTA resin (pre-equilibrated with 50 mM Tris-Cl, pH 7.5) for 1 hour, the resin was washed thrice with wash buffer (50 mM Tris-Cl, 150 mM NaCl, 10-50 mM imidazole, pH 7.5), and bound proteins were eluted by boiling the resin in 60 μl of 2× SDS-PAGE sample buffer for 15 minutes at 99°C. The eluates were processed for mass spectrometry as described below in the mass spectrometry section.

### Identification of PfNEDD8-conjugates in the parasite

Asynchronous cultures of HN8-expressing and wild type parasites at 10-15% parasitemia were harvested and processed for the isolation of parasites as described in the parasite culture section. The parasite pellets were resuspended in 10× pellet volume of the lysis buffer (100 mM Tris-Cl, 150 mm NaCl, 0.5 mM EDTA, 0.5% TritonX-100, pH 7.5, protease inhibitor cocktail (MERCK, 11873580001)), subjected to 2 cycles of lysis using the Diagenode Bioruptor^®^ (each cycle involved 30 seconds ON and 30 seconds OFF pulses for 5 minutes). The samples were centrifuged at 25000g for 30 min at 4°C, supernatants were transferred into a fresh micro centrifuge tube. The pellet was re-extracted with 3× pellet volume of the lysis buffer as described above, and the supernatant was pooled with the first supernatant. The protein amount in the supernatant was estimated using BCA method. The supernatant containing about 1.5 mg protein was incubated with rabbit anti-HA antibodies (10 μl slurry/mg protein) for overnight at 4°C with gentle shaking. The suspension was incubated with preequilibrated protein A/G magnetic beads (Thermo Fisher Scientific; 20 μl slurry/mg protein) for 1 hour at 4°C with gentle shaking. The flow through was separated and the beads were washed three times, each time with 1 ml of wash buffer (100 mM Tris-Cl, 150 mm NaCl, 0.5 mM EDTA, pH 7.5; protease inhibitor cocktails). The beads were resuspended in 100 μl of 2× SDS-PAGE sample buffer, boiled for 15 min, centrifuged and the supernatant was collected as eluate. A 20 μl aliquot of the eluate was assessed for the presence of HN8 along with appropriate controls (input, eluate, flow through, washes, and beads after elution) by western blotting using mouse anti-HA antibodies, followed by appropriate secondary antibodies as described in the western blotting section. The remaining 80 μl eluate was processed for mass spectrometry.

### Mass spectrometry

The eluates from immunoprecipitation and in-vitro neddylation assay were run on a 12% SDS-PAGE gel until the protein marker completely entered into the resolving gel. The gel was stained with coomassie blue and destained. The gel slice containing the protein band was excised, washed with 800 μl of 50 mM ammonium bicarbonate-acetonitrile solution (7:3 v/v), followed by sequential washing with 300-800 μl of 50 mM Ammonium bicarbonate and 800 μl of acetonitrile. The gel piece was vacuum dried, resuspended in 200 μl of 10 mM DTT for 45 min at 56°C, and incubated in 200 μl of 50 mM ammonium bicarbonate-55 mM iodoacetamide solution for 30 min at room temperature. The gel piece was washed with 700 μl of 50 mM ammonium bicarbonate, followed by with 700 μl of acetonitrile for 10 min. The gel piece was vacuum dried and treated with trypsin (15 ng/μl in 25 mM ammonium bicarbonate, 1 mM CaCl_2_; Promega or Roche sequencing grade) at 37°C for 16 hours. The peptides were extracted with 5% formic acid-30% acetonitrile solution, the extract was vacuum dried, dissolved in 20 μl of 0.1% TFA, desalted using C18 Ziptips (Merck, ZTC18M960), and eluted with 40 μl of 50% acetonitrile-5% formic acid solution. The eluate was vacuum dried and resuspended in 11 μl of 2% formic acid. 10 μl of the sample was run on the Q Exactive HF (Thermo Fischer Scientific) to perform HCD mode fragmentation and LC-MS/MS analysis. Raw data files were imported into the proteome discoverer v2.2 (Thermo Fischer Scientific), analysed and searched against the UniProt databases of *P. falciparum* 3D7 using the HT Sequest algorithm. The analysis parameters included enzyme specificity for trypsin, maximum two missed cleavages, carbidomethylation of cysteine, oxidation of methionine, deamidation of asparagine/glutamine as variable modifications. The precursor tolerance was set to 5 ppm and fragmentation tolerance was set to 0.05 Da. The peptide spectral matches (PSM) and peptide identifications grouped into proteins were validated using Percolator algorithm of proteome discoverer and filtered to 1% FDR. The protein hits from the HN8 immunoprecipitate were compared with those from the wild type *P. falciparum*. Protein hits identified in test eluate sample were compared with those in the control sample (without ATP). Common proteins or proteins with a difference of at least 5 times lower peptide spectrum matches (PSMs) in the controls compared to the test samples were excluded, and proteins common to at least three biological replicates with minimum 1 unique peptide were considered.

### Generation of *S. cerevisiae* Rub1 and Ubc12 knockout strains

*S. cerevisiae* NEDD8 is known as Related to ubiquitin 1(Rub1) and NEDD8 E2 is known as Ubc12. The Rub1 and Ubc12 genes were individually knocked out in BY4741 strain by replacement of the respective ORF with kanamycin expression cassette as has been described earlier [78]. For Rub1, the kanamycin cassette was amplified from pFA6a–kanMX4 plasmid using ScN8KO-F/ScN8KO-R primers. For Ubc12, the kanamycin cassette was PCR amplified from pFA6a–kanMX4 plasmid using ScU12KO-F/ScU12KO-R primers. The PCR products were gel extracted, and the purified PCR products were transformed into the BY4741 cells by lithium acetate method [79]. Briefly, the BY4741 strain was grown in 5 ml of complete YPD medium at 25°C and 250 rpm till OD_600_ of 1.0. The culture was centrifuged at 3000g for 5 minutes at 25°C, the pellet was washed twice with autoclaved milliQ™ water, and resuspended in 360 μl of transformation mix (100 mM lithium acetate, 34.7% PEG and 5 μg of calf thymus DNA) containing 1 μg of desired PCR product. The transformation reaction was vortexed for 30 seconds, incubated at room temperature for 30 minutes, incubated at 42°C for 15 minutes, followed by incubation at room temperature for 15 minutes. The transformed reaction was centrifuged at 3300g for 2 minutes at 25 °C, the cell pellet was resuspended in 100 μl nuclease free water, spread on YPD agarose plates containing G418 (350 μg/ml), and incubated at 25°C till resistant colonies appeared.

### Confirmation of knockouts and assessment of the growth rates of knockout strains

The wild type BY4741 strain and G418 resistant colonies were grown overnight in 5 ml of YPD medium at 25°C and 250 rpm. The cultures were harvested at 3000g for 5 minutes at 25°C, and the cell pellets were used for gDNA isolation by bead lysis method [80]. Briefly, the cell pellet was mixed with 0.3 g of glass beads and 300 μl of lysis buffer (10 mM Tris pH 8, 100 mM NaCl, 1 mM EDTA, 2% Triton X-100 and 1% SDS). The lysate was extracted with equal volume of phenol-chloroform mixture (1:1), centrifuged at 15000g for 5 minutes. The aqueous layer was transferred to a fresh tube and genomic DNA was precipitated using 1 ml of absolute ethanol. The DNA pellet was washed with 1 ml of 70% ethanol, air dried, and resuspended in 100 μl of nuclease free water. The gDNAs from wild type and knockout lines were used in PCR with primers specific for wild type locus (*rub1*Δ: ScN8-Fcon/ScN8-Rcon; *ubc12*Δ: ScU12-Fcon/ScU12-Rcon), 5’ integration (*rub1*Δ: ScN8-Fcon/kan2; *ubc12*Δ: ScU12-Fcon/kan2) and 3’ integration (*rub1*Δ: kan3/ScN8-Rcon; *ubc12*Δ: kan3/ScU12-Rcon). Primers specific for ScAtg18 (ScAtg18-F/ScAtg18-R) were used as a positive control.

For growth analysis, colonies of wild type, *rub1*Δ and *ubc12*Δ strains were inoculated in 5 ml YPD broth (with 350 μg/ml G418 for knockouts) and grown overnight at 25°C and 250 rpm. The overnight cultures were subcultured in triplicates with starting OD of 0.1, and their growth was monitored by measuring OD_600_ at every two hours for 14 hours. The OD values were plotted against time using the GraphPad Prism.

### Complementation of *rub1*Δ by PfNEDD8 and ScNEDD8

The PfNEDD8 coding sequence was amplified from pSMO-PfNEDD8 plasmid using primers PfN8-FScepi/PfN8-Rsp, and cloned at EcoRI/BamHI site of the pLE124 plasmid to obtain pLE124-PfN8, which would express it as an N-terminal 2xHA-tagged protein (HA-PfN8) under Cu^++^ inducible CUP1 promoter. This plasmid also contains *LEU2* marker for selection of transformed cells on minimal synthetic media without Leu. As a positive control, *S. cerevisiae rub1* (ScRub1) was also amplified from *S. cerevisiae* gDNA using the primers ScN8-Fepi/ScN8-Repi, and cloned at EcoRI/BamHI site of the pLE124 plasmid to obtain pLE124-ScN8, which would express it as an N-terminal 2xHA-tagged protein (HA-ScN8) under Cu^++^ inducible CUP1 promoter. The *rub1*Δ and *ubc12*Δ cells were transformed with pLE124-PfN8 or pLE124-ScN8 as described above, and the transformants were selected on synthetic media without Leu to obtain complemented cells.

Expression of HA-PfN8 and HA-*rub1* was checked in complemented strains by western blotting using anti-HA antibodies. Briefly, BY4741 was grown in YPD medium; knock-out strains *rub1*Δ and *ubc12*Δ were grown in YPD media with G418 (350 μg/ml); the complemented strains *rub1*Δ [HA-PfN8], *rub1*Δ [HA-*rub1*], *ubc12*Δ[HA-PfN8] and *ubc12*Δ[HA-*rub1*] were grown in synthetic media without Leu, induced with 100 μM CuSO4. All the cultures were grown to exponential phase and harvested at 3000g for 5 minutes at 25°C to obtain cell pellets and processed as mentioned before [81]. The cell pellets corresponding to 5OD_600_ were resuspended in 250 μl of 1.85 M NaOH, incubated on ice for 10 minutes, 250 μl of 50% trichloroacetic acid was added to the suspension, and it was centrifuged at 18000g for 5 minutes at 4°C. The supernatant was aspirated and the pellet was washed with 1 ml of 1 M Tris pH 7.5, resuspended in 120 μl of 2× SDS-PAGE sample buffer, boiled for 10 minutes, centrifuged at 18000g for 30 minutes, and supernatant was transferred to a fresh micro centrifuge tube. Identical volumes of supernatants were processed for western blotting using rabbit anti-HA antibody, followed by HRP-conjugated goat anti-rabbit antibody as described in the western blotting section. For loading control, the blot was stripped using 0.2 N NaOH, and developed using mouse anti-phosphoglycerate kinase (PGK) antibodies, followed by appropriate secondary antibody as described in the western blot section.

### Immunoprecipitation of HA-PfN8 and HA-*rub1*

Wild type, *rub1*Δ[HA-PfN8] and *rub1*Δ[HA-rub1] strains were grown in YPD broth for overnight at 25°C and 250 rpm. The cultures were harvested at 3000g for 5 minutes at 25°C. Cell pellets corresponding to 60OD_600_ were resuspended in 360 μl of ice-cold lysis buffer (10 mM Tris, 10 mM HEPES, 150 mM NaCl, 0.5 mM EDTA, 0.5% NP-40, pH 7.5 and 1× protease inhibitor cocktail), incubated in ice for 15 min, mixed with 0.5 g of glass beads, and the cells were disrupted by bead beating for 80 seconds. The samples were incubated in ice for 2 minutes and again subjected to bead beating two times. The sample was incubated in ice for 10 minutes, centrifuged at 25000g for 20 minutes at 4°C, and the supernatant was transferred to a micro centrifuge tube. The pellet was re-extracted with 240 μl of the lysis buffer, and the supernatant was pooled with the first supernatant. The protein amount in the supernatant was estimated using BCA method.

10 μg of mouse anti-HA antibody (Thermo Fisher Scientific) was coupled with 10 μl of protein A/G magnetic beads pre-equilibrated with wash buffer (10 mM Tris, 10 mM HEPES, 150 mM NaCl, 0.5 mM EDTA, 0.1% NP-40, pH 7.5) for 2 hours at 4°C in a tube rotator. 2 mg of the protein extract was added to the anti-HA antibody-coupled beads, incubated overnight at 4°C, washed with wash buffer five times, and bound proteins were eluted by boiling the beads in 100 μl of 2× SDS-PAGE sample buffer for 10 min. A 20 μl aliquot of the eluate was processed for western blotting along with appropriate controls (input, flow through and washes) using rabbit anti-HA antibody, followed by HRP-conjugated goat anti-rabbit antibody as described in the western blotting section. The remaining 80 μl of the eluate was processed for mass spectrometry as described in the mass spectrometry section.

## RESULTS AND DISCUSSION

### Malaria parasites encode two types of NEDD8 proteins

Searches of the *Plasmodium* genome databases using human and *S. cerevisiae* NEDD8 sequences identified several sequences, which were analyzed for the presence of conserved motifs and amino acid residues unique to the characterized NEDD8 proteins of model organisms [26,82,83]. The top 3 hits (XP_001350369.1, e-value = 7e-29; XP_001350526.1, evalue =3e-26; XP_001349866.1, e-value= 3.6e-24) were used in reverse BLAST of the UniProt database, which revealed that XP_001350369.1 and XP_001350526.1 were closer to ubiquitin than NEDD8, hence, these were excluded in further analysis. XP_001349866.1 is likely to be a *P. falciparum* NEDD8 homolog (PfNEDD8), and it showed y 46%-52% sequence identity with NEDD8 proteins of model organisms (Fig. 1A). Like ubiquitin, NEDD8 is synthesized as a precursor that requires processing of its C-terminal tail to free the 76^th^ Gly residue at the C-terminus for conjugation. Interestingly, PfNEDD8 contains 76 amino acid residues only, with Gly as the 76^th^ residue, suggesting that it is readily available for conjugation (Fig. 1A). Human and *P. falciparum* ubiquitins are 58% and 51% identical to their respective NEDD8 proteins. As expected for ubiquitin-family proteins, human NEDD8 and ubiquitin have similar secondary structure [82]. Amino acid residues at 31^st^, 32^nd^ and 72^nd^ positions are unique to ubiquitin and NEDD8 [32], which have been demonstrated to be critical in discrimination of ubiquitin and NEDD8 by their respective enzymatic machinery. For example, ubiquitin has an Arg residue in the 72^nd^ position, whereas NEDD8 contains a non-positively charged residue in the same position, which dictate specific activation by their respective E1 enzymes [84]. The 72^nd^ residue in PfNEDD8 and other *Plasmodium* NEDD8 proteins is Gln, whereas *Plasmodium* ubiquitin has Arg at the corresponding position. Residues 31^st^ and 32^nd^ in all characterized NEDD8 are Glu-Glu, which have been shown to be pivotal for cullin-1 neddylation. Interestingly, substituting the corresponding residues of ubiquitin with Glu-Glu caused ubiquitylation of cullin-1 [32].

**Fig. 1.**
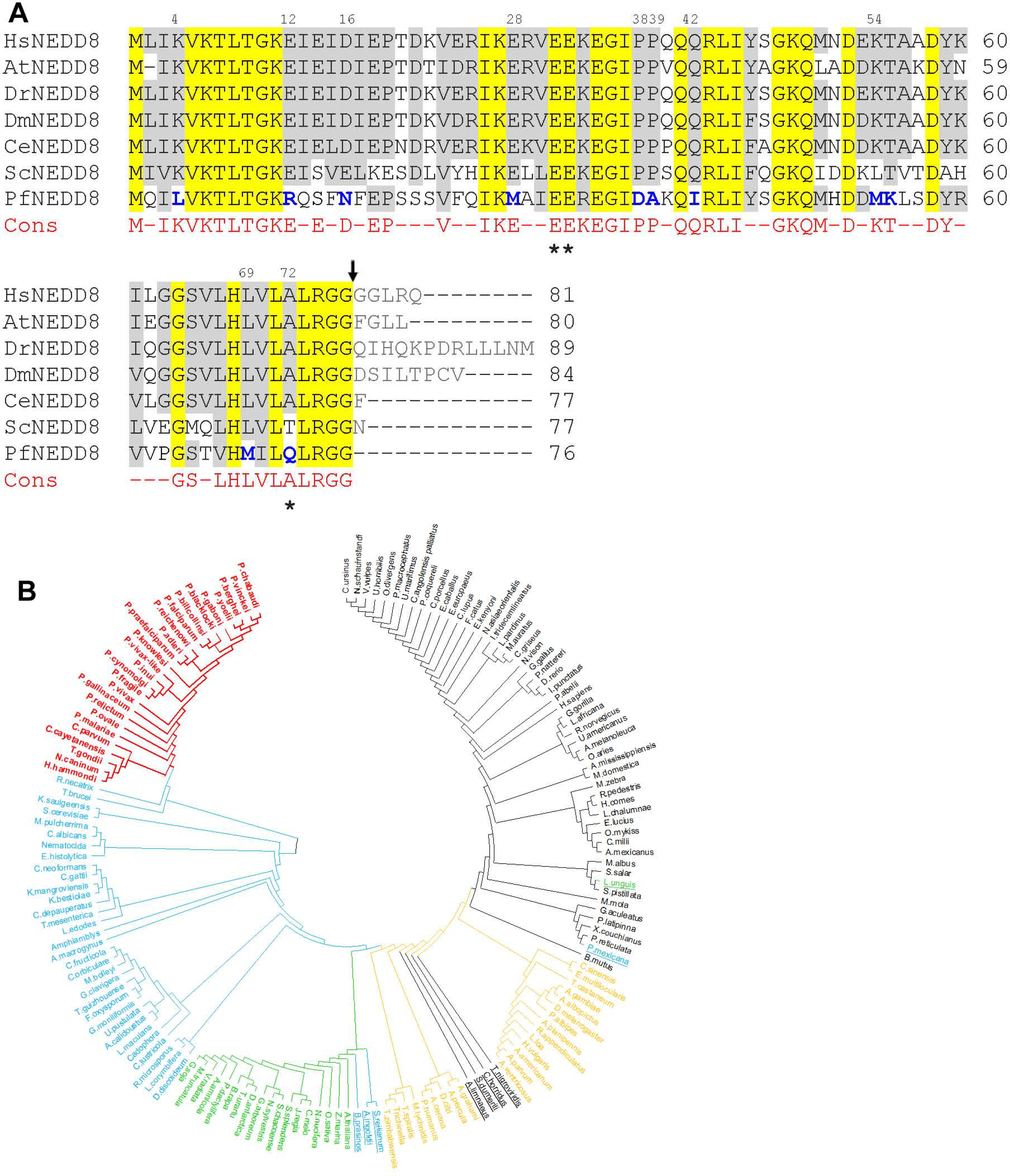
**A. Sequence alignment of PfNEDD8.** The amino acid sequences of 160 NEDD8 proteins from various organisms representing protozoans, metazoans, plants and fungi were aligned. Shown is the alignment of *P. falciparum* NEDD8 (PfNEDD8) with homologs from the indicated model organisms (*H. sapiens*: HsNEDD8, *A. thaliana*: AtNEDD8, *D. rerio*: DrNEDD8, *D. melanogaster*: DmNEDD8, *C. elegans*: CeNEDD8, *S. cerevisiae*: ScNEDD8). Also shown is the consensus (Cons) sequence in at least 75% of the 160 NEDD8 proteins, with hyphens representing variation at these positions. The conserved residues are yellow shaded, and grey shaded are physicochemically similar residues in at least 5 of the 7 sequences. The PfNEDD8 amino acid residues that are different from the consensus in physicochemical properties are in bold blue font. Amino acid residues marked with asterisks differentiate ubiquitin and NEDD8 for being correctly recognized by their respective enzymatic machinery. Numbers on the top of the sequence indicate position of that amino acid residue in PfNEDD8. The black arrow represents the processing site to remove the C-terminal tail. **B. Phylogenetic analysis of NEDD8 homologs**. 160 NEDD8 protein sequences of the indicated organisms representing metazoans, plants, fungi and protozoans were aligned using the CLUSTAL W program, and subjected to phylogenetic analysis using the maximum likelihood method. The NEDD8 homologs of apicomplexans are in red, non-apicomplexan protists and fungi are in blue, plants are in green, invertebrates are in yellow and vertebrates are in black. The outlier NEDD8 homologs are underlined.

NEDD8 is highly conserved across *Plasmodium* species (>89% identity). However, sequence alignment of *Plasmodium* NEDD8 proteins revealed two forms of NEDD8 (Fig. S1). The majority of human and primate malaria parasite NEDD8 proteins lack C-terminal tails, whereas the NEDD8 proteins of rodent and avian malaria parasites contain C-terminal tails. The NEDD8 proteins of human malaria parasites *P. ovale* and *P. malariae* have C-terminal tails. The presence of C-terminal tail in NEDD8 proteins of rodent and other malaria parasites suggests requirement for processing by a C-terminal hydrolase, which may represent an additional layer of regulation compared to the *Plasmodium* species containing tail-less NEDD8 proteins, which would bypass the processing step. We compared *Plasmodium* NEDD8 proteins with homologs from a diverse range of eukaryotes representing metazoans, plants, fungi and protozoans. Notably, at 11 amino acid residues in the *Plasmodium* NEDD8 proteins are strikingly different in physicochemical properties from those present in the majority of NEDD8 homologs at corresponding positions (Fig. S1), which may represent specificity for *Plasmodium* neddylation pathway enzymes. Many of these residues are also conserved in other apicomplexan NEDD8 homologs. One such notable feature is the presence of Gln at the 72^nd^ position in apicomplexan NEDD8 homologs, the key specificity determinant of NEDD8 for NAE, whereas Ala/Thr occupies the corresponding position in other NEDD8 proteins analyzed in this study. The residues 31^st^ and 32^nd^ in NEDD8 homologs of some apicomplexan parasites like *T. gondii* and *C. parvum* are Gln-Glu as opposed to Glu-Glu in the majority of NEDD8 proteins analyzed, including Plasmodium. Given the importance of two residues for cullin neddylation, it may indicate organism specific tuning for neddylation. As mentioned above, certain *Plasmodium* species among all apicomplexan parasites did not have the C-terminal tail. Among the *non-Plasmodium* NEDD8 homologs, only *Nematocida, H. vulgaris* and *Amphiamblys* encode NEDD8 proteins without C-terminal tails. Interestingly, the Atg8 proteins of *Plasmodium* parasites also lacks C-terminal tail [19]. It is required to investigate whether tail-less NEDD8 and Atg8 offer any advantage. Consistent with the conservation of several unique amino acid residues, all apicomplexan NEDD8 proteins share a common node in the phylogenetic tree (Fig. S1B). The apicomplexan NEDD8 proteins are more closely related to the NEDD8 proteins of other unicellular protists and fungi compared to those of other organisms, as they have a common ancestor. Baring few outliers, the plant, vertebrate and invertebrate NEDD8 proteins came together with the respective group of organisms in the phylogenetic tree.

### Recombinant PfNEDD8 gets conjugated in an ATP-dependent manner

To experimentally validate if the predicted PfNEDD8 is a functional protein, we produced it as a C-terminal fusion of His-tagged SUMO (SN8) using *E. coli* expression system (Fig. S2). Since NAE and Ubc12 homologs of *Plasmodium* remain to be identified and characterized, we used *P. falciparum* erythrocytic stage parasite lysate as a source of neddylating enzymes and substrates for in vitro neddylation experiment in the presence or absence of ATP. The western blot of ATP-containing reaction samples showed the presence of higher molecular weight bands, which were absent in the reaction samples without ATP, indicating that SN8 was conjugated to parasite proteins in an ATP-dependent manner (Fig. 2). This is consistent with previous reports of ATP-dependent conjugation of ULMs [85,86]. To identify SN8-conjugates, we purified the reaction samples and the eluates were subjected to mass spectrometric analysis. Several proteins were identified in ATP-containing reaction, which were absent in the reaction samples without ATP (Table 1). The important candidates are two putative cullins and the suppressor of kinetochore protein 1 (Skp-1). Since cullins are the bonafide neddylation substrates, the conjugation of SN8 to cullins indicate that the predicted PfNEDD8 is a true NEDD8 and *Plasmodium* has a functional neddylation pathway. Skp-1 is an indispensable component of the Skp1-Cullin-F-box protein E3 ubiquitin ligase complex (SCF) in which it serves as an adapter protein that links F-box proteins to cullin-1 [87]. In addition to cullin and Skp-1, SCF also contains Rbx-1 and F-box proteins. Rbx-1 is a RING-type zinc finger domain protein that recruits charged E2-ubiquitin to SCF complex. Rbx-1 has also been demonstrated to function as an E3 for neddylation of cullin in humans [30,88]. F-box proteins are responsible for recruitment of the substrate to be ubiquitylated. However, *P. falciparum* Rbx-1 and F-box proteins were not present in the mass spectrometry analysis, which could be due to dissociation of the SCF complex under our experimental conditions. In addition to cullins, phosphatidylinositol 3-kinase, ER lumen protein retaining receptor 1, and coatomer subunit delta were also identified in the mass spectrometry analysis. These molecules have not been reported to be neddylation substrates yet. It is possible that these proteins are associated with cullins or the neddylation pathway. We could not identify homologs of *P. falciparum* neddylating enzymes, which might be due to transient nature of neddylation reactions.

**Fig. 2.**
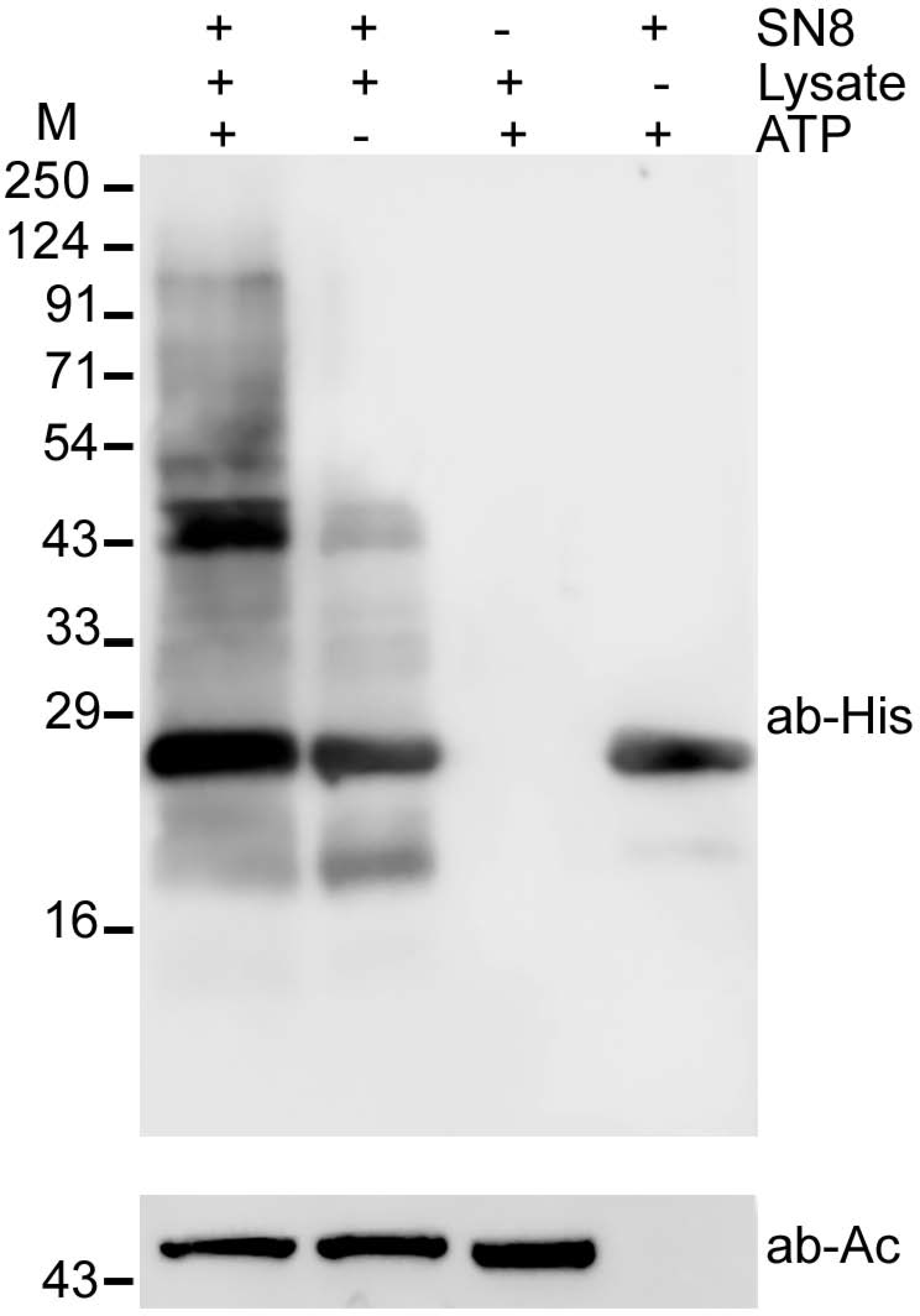
Western blot analysis of in vitro neddylation. Conjugation of recombinant SUMO-PfNEDD8 (SN8) was carried out using parasite lysate (Lys) and ATP at 37°C for 1 hour. The reactions were stopped and subjected to immunoblotting using anti-His antibody. The presence (+) or absence of reaction component is indicated. β-actin was used as a loading control for reactions containing the parasite lysate. The positions of proteins size markers are shown in kDa (M).

**Table 1:** Proteins identified in the in-vitro neddylation assay: The proteins listed are exclusive to ATP-containing reaction and common in at least 2 of the three independent experiments. Shown above are the score (Sc), coverage (Co), and number of unique peptides (UP) for each protein. The predicted role is based on experimentally validated homologs in other systems.

| ID | Protein | Experiment 1 |  |  | Experiment 2 |  |  | Experiment 3 |  |  |
| --- | --- | --- | --- | --- | --- | --- | --- | --- | --- | --- |
|  |  | Sc | Co | UP | Sc | Co | UP | Sc | Co | UP |
| Q8IAU5<br>(PF3D7_0811000) | Cullin-1 | 43.2 | 26.3 | 15.0 | 15.7 | 15.6 | 6.0 | 32.3 | 17.3 | 10.0 |
| C6KTD3<br>(PF3D7_0629800) | Cullin-like protein | 7.1 | 11.3 | 2.0 | 17.8 | 14.5 | 6.0 | - | - | - |
| Q8I3V5<br>(PF3D7_0515300) | Phosphatidylinositol 3-kinase | 11.5 | 11.1 | 3.0 | 61.1 | 11.3 | 12.0 | - | - | - |
| Q8ID38<br>(PF3D7_1367000) | Suppressor of kinetochore protein 1 | 7.0 | 14.8 | 2.0 | - | - | - | 5.9 | 14.8 | 2.0 |
| A0A143ZVV1<br>(PF3D7_1330400.2) | ER lumen protein retaining receptor 1 | 5.2 | 18.8 | 2.0 | - | - | - | 4.4 | 14.3 | 2.0 |
| Q8II16<br>(PF11_0359) | Coatomer subunit delta | 8.0 | 8.8 | 4.0 | - | - | - | 1.8 | 6.0 | 1.0 |

### PfNEDD8 complemented *S. cerevisiae* Rub1

As neddylation pathway is conserved across eukaryotes, we investigated if PfNEDD8 complements *S. cerevisiae* NEDD8 homolog, Rub1. First, we individually knocked-out rub1 and ubc12 in *S. cerevisiae* wild type strain BY4741, and PCR of the genomic DNAs of mutants (*rub1*Δ and *ubc12*Δ) confirmed the replacement of target genes by kanamycin cassette (Fig. 3A and 3B). The growth rates of mutants were similar to that of wild type strain (Fig. S3), which is in agreement with a previous report of showing dispensability of neddylation pathway for normal growth of *S. cerevisiae* [89]. Next, in the mutants, we episomally expressed HA-tagged PfNEDD8 (*rub1*Δ[HA-PfN8] and *ubc12*Δ[HA-PfN8]) or HA-tagged rub1 (*rub1*Δ [HA-*rub1*] and *ubc12*Δ[HA-*rub1*]), and confirmed the expression of HA-tagged proteins by western blot (Fig. 3C). The blots of *rub1*Δ-complemented strains (*rub1*Δ[HA-PfN8] and *rub1*Δ[HA-*rub1*]) had a prominent band above the 91 kDa marker, which was absent in the wild type, knockouts (*ub1*Δ and *ubc12*Δ) and the *ubc12*Δ-complemented (*ubc12*Δ[HA-PfN8] and *ubc12*Δ[HA-*rub1*]) strains, indicating that conjugation of both PfNEDD8 and Rub1 involved the neddylation pathway. The prominent bands could be *S. cerevisiae* cullins (cdc53: 93.95 kDa, CUL3: 87.1 kDa). Several other bands were also observed in the *rub1*Δ-complemented strains only, which may be conjugates of Rub1/NEDD8 in addition to cullins.

**Fig. 3.**
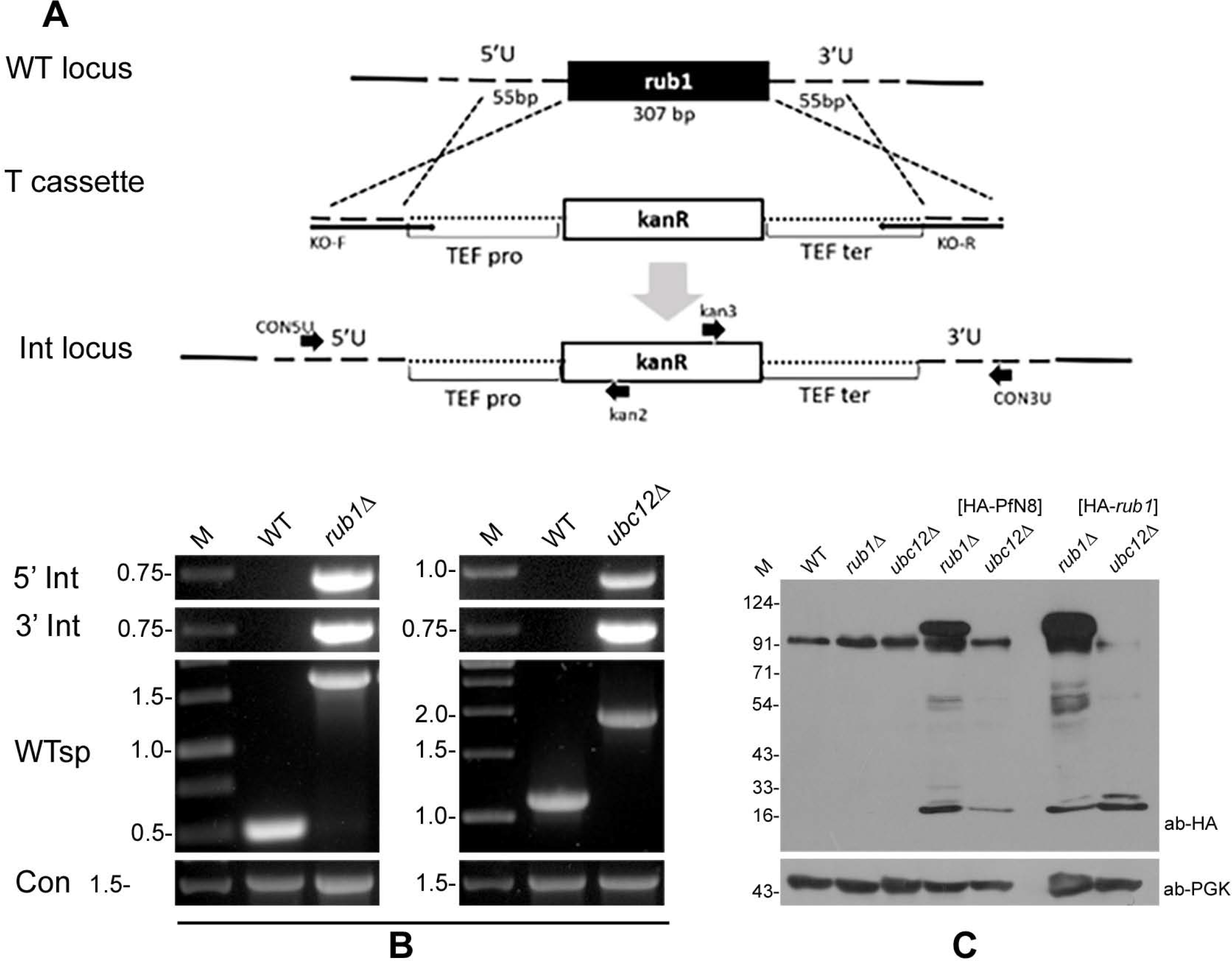
Complementation of ScRub1 by PfNEDD8. **A. Schematic for generation of *rub1Δ* or *ubc12Δ*.** The wild type locus (WT locus) was replaced with a linear kanamycin cassette flanked by the homology regions of the target locus (T cassette) via a double crossover homologous recombination, resulting in the integration locus (Int locus). The kanamycin coding sequence (kanR) is under the control of translation elongation factor promoter (TEF pro) and terminator (TEF ter). The arrows indicate the primer positions. **B. Confirmation of knockout.** The knockout was confirmed by PCR of the gDNAs of wild type (WT), *rub1Δ* and *ubc12Δ* strains using locus-specific primers (5’ Int for 5’ integration locus, 3’ Int for 3’ integration locus, WTsp for wild type locus and Con for ScAtg18). The ethidium bromide stained agarose gel shows PCR products, with DNA markers in kbp (M). **C. Western blot analysis of complemented strains.** HA-PfNEDD8 (HA-PfN8) or HA-ScRub1 (HA-rub1) were episomally expressed in *rub1Δ* and *ubc12Δ* strains. The lysates of wild type (WT), *rub1Δ, ubc12Δ*, HA-PfNEDD8-complemented (HA-PfN8), and HA-ScNEDD8-complemented (HA-rub1) *rub1Δ* or *ubc12Δ* strains were processed for western blotting using anti-HA antibodies (ab-HA). The lanes containing lysates of complemented strains show a free NEDD8/Rub1 protein, whereas the lanes with *rub1Δ* [HA-PfN8] and *rub1Δ* [HA-*rub1*] strains also contain a high molecular weight band (>91kDa) and some lower molecular weight conjugates, which are absent in other lanes. The phosphoglycerate kinase (anti-PGK) was used as a loading control. The protein size markers are in kDa (M).

We used the lysates of *rub1*Δ-complemented strains (*rub1*Δ [HA-PfN8] and *rub1*Δ [HA-*rub1*]) and wild type strain for immunoprecipitation using anti-HA antibody. The western blot confirmed the presence of free NEDD8/Rub1 and its conjugates of different sizes, which were present in the eluates from *rub1*Δ-complemented strains only (Fig. 4). Mass spectrometry of the eluates from *rub1*Δ-complemented strains identified five unique proteins (Tables 2, 3 and 4), including cdc53. The other relevant proteins included NEDD8 activating enzyme catalytic subunit Uba3, NEDD8 activating enzyme regulatory subunit Ula1 and SCF subunits. Although cdc53 was identified in the *rub1*Δ[HA-PfN8] immunoprecipitate in three independent experiments, the reproducibility of neddylation enzymes was less as compared to that of *rub1*Δ[HA-*rub1*] immunoprecipitate. This could be due to different enzyme kinetics of PfNEDD8 and ScRub1 conjugation with the substrates, which may be attributed to differences in the efficacy of *S. cerevisiae* NAE and Ubc12 to conjugate PfNEDD8 compared to Rub1. It may be interesting to reiterate here that PfNEDD8 differs from the majority of NEDD8 proteins at 11 positions, and some of these residues might play important roles in determining neddylation efficiency and specificity of PfNEDD8. This also suggests that subtle differences in NEDD8 bring about organism-specific tuning of neddylation pathway. Taken together, conjugation of PfNEDD8 to *S. cerevisiae* cdc53 demonstrates that PfNEDD8 is a genuine NEDD8 of *P. falciparum*. This also highlights functional conservation of NEDD8 between *Plasmodium* and *S. cerevisiae*, and provides the proof of concept for the utility of *S. cerevisiae* to study *Plasmodium* NEDD8.

**Fig. 4.**
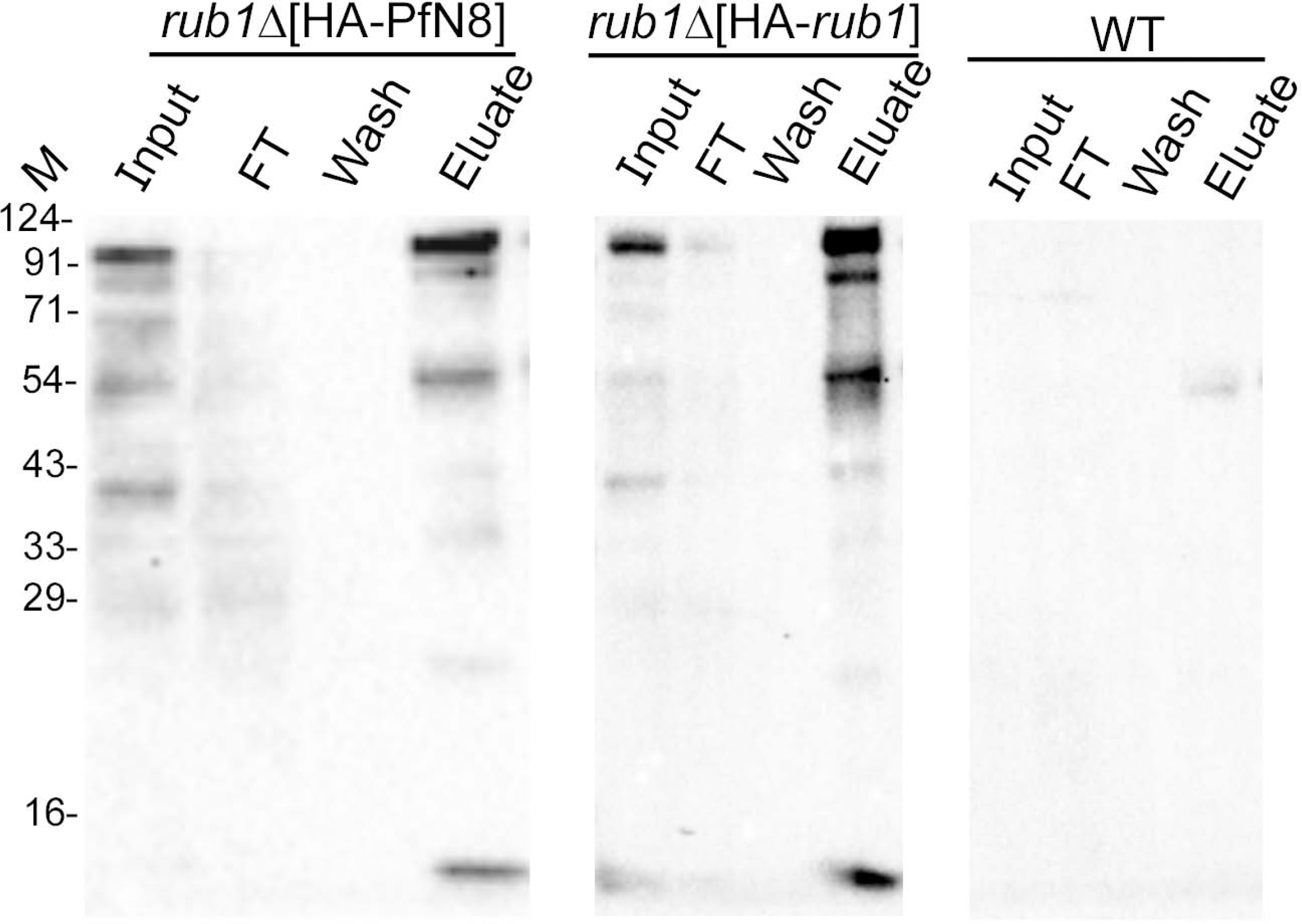
Immunoprecipitation of PfNEDD8 and Rub1 from complemented strains. The lysates of *rub1*Δ [HA-PfN8], *rub1*Δ [HA-*rub1*] and wild type (WT) strains were immunoprecipitated with mouse anti-HA antibody. The eluates (Eluate) along with the extract (Input), flow through (FT) and wash (Wash) were analyzed by western blotting using rabbit anti-HA antibodies. The eluate lanes contain the respective target protein, indicating successful immunoprecipitation. The sizes of protein size markers are in kDa (M).

**Table 2.**
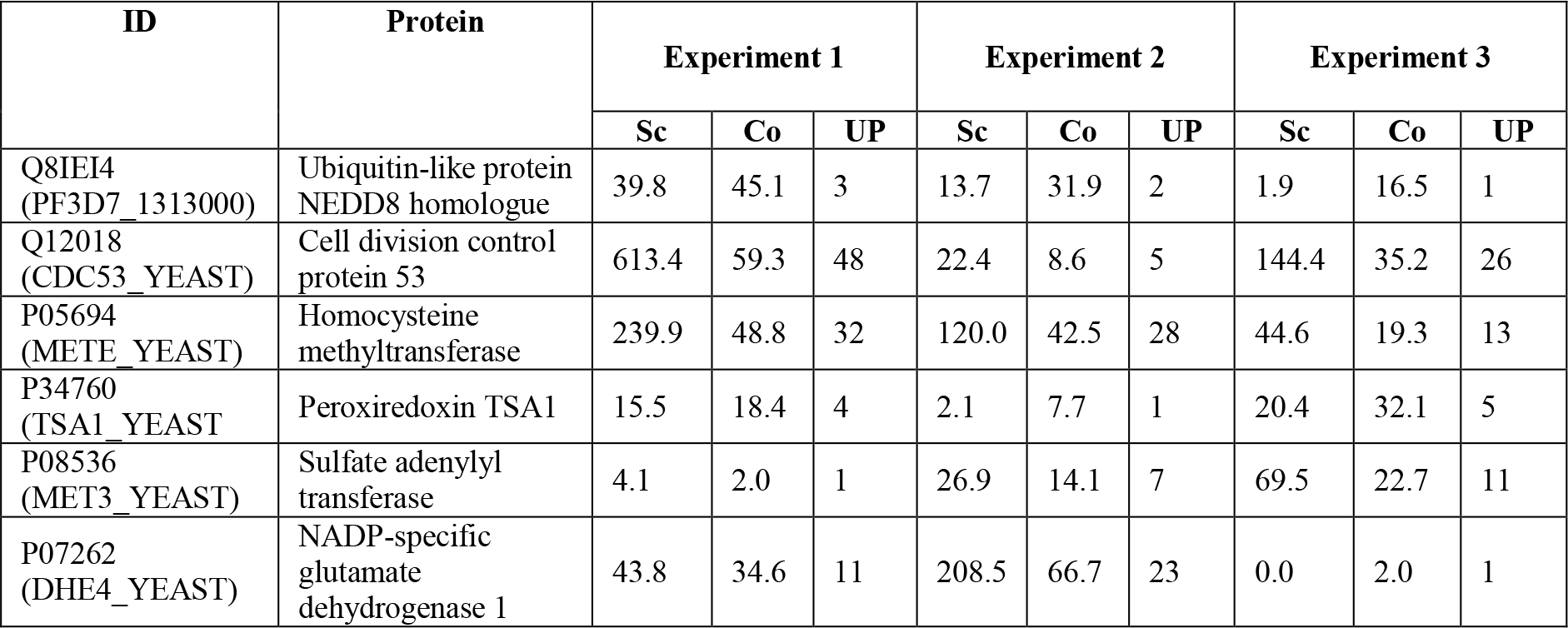
Proteins identified in the immunoprecipitate of *rub1*Δ[HA-PfN8]. Shown are score (Sc), coverage (Co) and unique peptides (UP) for each protein.

| ID | Protein | Experiment 1 |  |  | Experiment 2 |  |  | Experiment 3 |  |  |
| --- | --- | --- | --- | --- | --- | --- | --- | --- | --- | --- |
|  |  | Sc | Co | UP | Sc | Co | UP | Sc | Co | UP |
| Q8IEI4<br>(PF3D7_1313000) | Ubiquitin-like protein<br>NEDD8 homologue | 39.8 | 45.1 | 3 | 13.7 | 31.9 | 2 | 1.9 | 16.5 | 1 |
| Q12018<br>(CDC53_YEAST) | Cell division control<br>protein 53 | 613.4 | 59.3 | 48 | 22.4 | 8.6 | 5 | 144.4 | 35.2 | 26 |
| P05694<br>(METE_YEAST) | Homocysteine<br>methyltransferase | 239.9 | 48.8 | 32 | 120.0 | 42.5 | 28 | 44.6 | 19.3 | 13 |
| P34760<br>(TSA1_YEAST) | Peroxiredoxin TSA1 | 15.5 | 18.4 | 4 | 2.1 | 7.7 | 1 | 20.4 | 32.1 | 5 |
| P08536<br>(MET3_YEAST) | Sulfate adenylyl<br>transferase | 4.1 | 2.0 | 1 | 26.9 | 14.1 | 7 | 69.5 | 22.7 | 11 |
| P07262<br>(DHE4_YEAST) | NADP-specific<br>glutamate<br>dehydrogenase 1 | 43.8 | 34.6 | 11 | 208.5 | 66.7 | 23 | 0.0 | 2.0 | 1 |

**Table 3.**
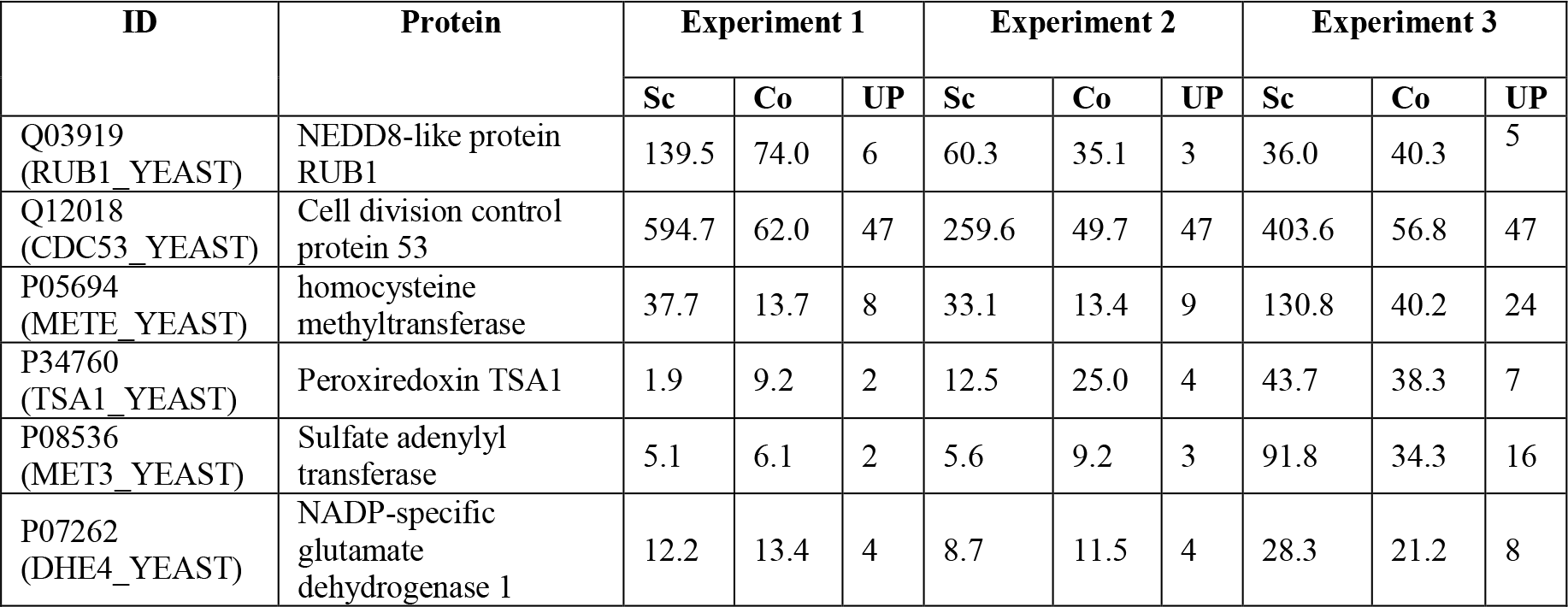
Proteins identified in the mass spectrometry of immunoprecipitate of *rub1*Δ[HA-*rub1*]. Shown are score (Sc), coverage (Co) and unique peptides (UP) for each protein.

**Table 4.** Neddylation pathway and substrate proteins common in the mass spectrometry of immunoprecipitates of *rub1*Δ[HA-*rub1*] (all three experiments) and *rub1*Δ[HA-PfN8] (at least once). Shown are score (Sc), coverage (Co) and unique peptides (UP) for each protein.

| ID | Protein | *rub1Δ* [HA-PfN8] |  |  | *rub1Δ* [HA-*rub1*] |  |  |  |  |  |  |  |  |
| --- | --- | --- | --- | --- | --- | --- | --- | --- | --- | --- | --- | --- | --- |
|  |  | Experiment 1 |  |  | Experiment 1 |  |  | Experiment 2 |  |  | Experiment 3 |  |  |
|  |  | Sc | Co | UP | Sc | Co | UP | Sc | Co | UP | Sc | Co | UP |
| P52286<br>(SKP1_YEAST) | Suppressor of kinetochore protein 1 | 18.1 | 32.5 | 6 | 17.2 | 23.7 | 5 | 3.9 | 26.3 | 3 | 38.3 | 28.9 | 6 |
| Q99344<br>(UBA3_YEAST) | NEDD8-activating enzyme E1 catalytic subunit | 4.2 | 3.7 | 1 | 17.0 | 14.0 | 4 | 0.0 | 5.7 | 2 | 8.7 | 5.0 | 1 |
| Q08273<br>(RBX1_YEAST) | RING-box protein HRT1 | 18.7 | 31.4 | 2 | 15.0 | 11.6 | 1 | 28.9 | 24.0 | 3 | 7.3 | 11.6 | 1 |
| P24814<br>(GRR1_YEAST) | SCF E3 ubiquitin ligase complex F-box protein GRR1 | 61.8 | 18.2 | 20 | 48.8 | 9.9 | 9 | 15.2 | 7.7 | 8 | 2.1 | 0.7 | 1 |
| Q12059<br>(ULA1_YEAST) | NEDD8-activating enzyme E1 regulatory subunit | 2.4 | 2.6 | 1 | 12.3 | 8.2 | 4 | 16.0 | 14.1 | 7 | 12.2 | 12.8 | 5 |
| P47005<br>(DAS1_YEAST) | F-box protein DAS1 | 71.7 | 31.7 | 19 | 15.8 | 6.8 | 4 | 0.0 | 6.3 | 3 | 25.3 | 12.5 | 9 |
| P38352<br>(SAF1_YEAST) | SCF-associated factor 1 | 48.0 | 26.7 | 13 | 6.8 | 5.0 | 3 | 0.0 | 11.8 | 5 | 127.7 | 43.3 | 19 |

### Parasite proteins undergo neddylation

To investigate if neddylation is operational in malaria parasites, we episomally expressed HA-tagged PfNEDD8 (HN8) in *P. falciparum*. Western blot of the wild type and HN8-expressing parasites with anti-HA antibodies showed both free (~12 kDa) and conjugated forms of PfNEDD8 in HN8-expressing parasites only (Fig. 5A), including two prominent bands around 91 kDa and 124 kDa.

**Fig. 5.**
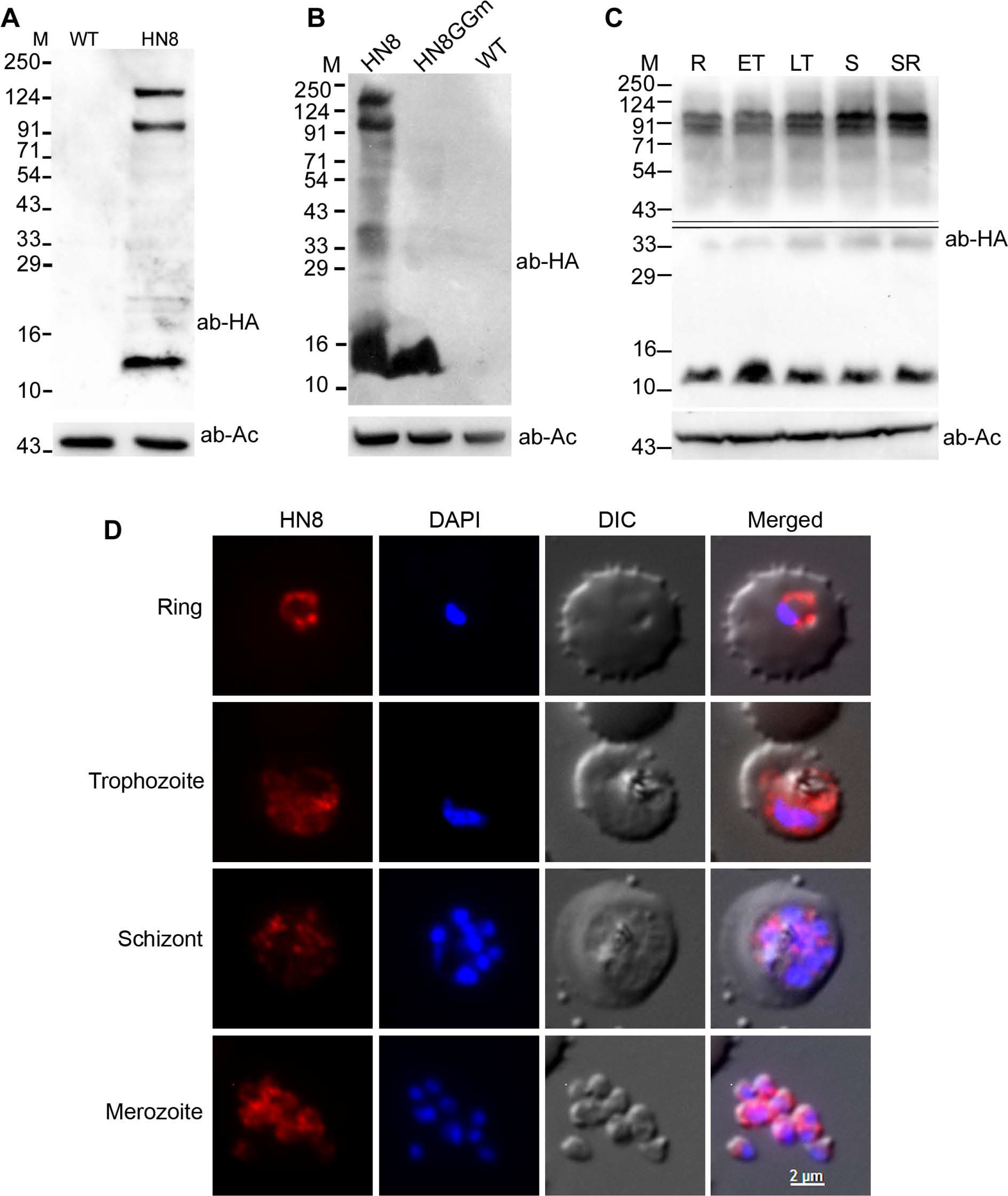
Expression, conjugation and localization of *P. falciparum* NEDD8. **A**. Lysates of the wild type (WT) and HA-PfNEDD8–expressing (HN8) *P. falciparum* parasites were processed for western blotting using anti-HA antibodies (ab-HA). β-actin was used as a loading control (ab-Ac). The blot shows a prominent band close to the predicted size of HA-PfHANEDD8 and two high molecular bands in the HN8 lane only. The protein markers are in kDa (M). **B.** The HΛ-PíNEDD8-expressing (HN8), mutant HA-PfNEDD8 (HN8GGm) and wild type (WT) parasites were processed for western blotting using anti-HA antibodies (ab-HA), and β-actin was used as a loading control (ab-Ac). Note that the prominent high molecular bands are present in HN8 lane only. Protein markers are in kDa (M). **C.** A synchronized culture of HA-PfNEDD8–expressing *P. falciparum* parasites was harvested at ring (R), early trophozoite (ET), late trophozoite (LT) and schizont/ring (SR) stages, and the parasites (1×10^8^ parasites/lane) were processed for western blotting using anti-HA antibodies (ab-HA). β-actin was used as a loading control (ab-Ac). The blot shows two prominent high molecular weight bands along with the free HA-NEDD8, indicating neddylation throughout the erythrocytic cycle. The protein size markers are in kDa (M) The double line in the top panel marks two different blots, which were developed separately to minimize over saturation of the blot due to the band around 12 kDa. **D.** Fixed HA-NEDD8-expressing *P. falciparum* parasites of the indicated stages were evaluated for localization of HA-NEDD8 using anti-HA antibodies. The images show HA-NEDD8 specific signal (HA-NEDD8), nucleic acid staining (DAPI), the parasite and the erythrocyte boundaries (DIC), and the merged of all three images (Merged). HA-NEDD8 signal is present throughout the parasite in all the stages shown, except the food vacuole. The scale bar shown is identical for all the images.

To confirm if the conjugation of PfNEDD8 is through its C-terminal Gly residue, we generated *P. falciparum* parasites episomally-expressing HA-tagged PfNEDD8 mutant (HN8GGm) that contains G75A/G76A substitution, which would render HN8GGm non-conjugatable. The western blot of HN8GGm-expressing parasites showed free HN8GGm but not any conjugated forms (Fig. 5B), confirming that PfNEDD8 undergoes conjugation through its C-terminus Gly residue and that the high molecular weight bands observed in the western blot of HN8-expressing parasites are indeed covalent conjugates of PfNEDD8. Western blot of different stages of HN8-expressing parasites showed PfNEDD8 conjugates, indicating that neddylation pathway is operational in these stages (Fig. 5C). Immunofluorescence assay of different stages of HN8-expressing parasites showed PfNEDD8 throughout the cytosol, except the food vacuole (Fig. 5D).

### The neddylation inhibitor MLN4924 did not affect neddylation in Plasmodium

Pevonedistat or MLN4924 is a general neddylation inhibitor, which has been shown to be a potent and cell-permeable inhibitor of NAE [90,91]. It is an AMP mimetic and preferentially binds the adenylate site of NAE that already has NEDD8 conjugated to its active site cysteine. In the MLN4924-NAE-NEDD8 complex, MLN4924 attacks the NEDD8, forming a sulfamate adduct, which renders NAE non-functional. It inhibits neddylation in mammalian cells at nanomolar concentration without affecting conjugation of other ULMs [91]. Given that neddylation pathway is implicated in cancer prognosis and progression [92,93], MLN4924 has also been shown to have potent anti-cancer activity in human/mouse cell culture models and pre-clinical studies [53,94–96]. MLN4924 has recently been shown to be toxic for *P. falciparum* erythrocytic stage development, and the anti-parasitic effect was attributed to inhibition of neddylation [61]. To check if the toxicity of MLN4924 is through inhibition of neddylation, we assessed *P. falciparum* erythrocytic stage development in the presence of MLN4924. The parasite growth was inhibited with the IC_50_ concentration of 14.4 ± 1.9 μM (Fig. 6A), which is over 2900-fold higher than the IC_50_ concentration reported for mammalian cells (4.7 nM) [90]. However, MLN4924 even at 12× IC_50_ concentration did not noticeably affect conjugation of PfNEDD8 (Fig. 6B), indicating that it inhibits parasite growth by targeting some other process. This also suggests that *Plasmodium* NAE is significantly different from its human homolog and it should be possible to make specific inhibitors of *Plasmodium* NAE.

**Fig. 6.**
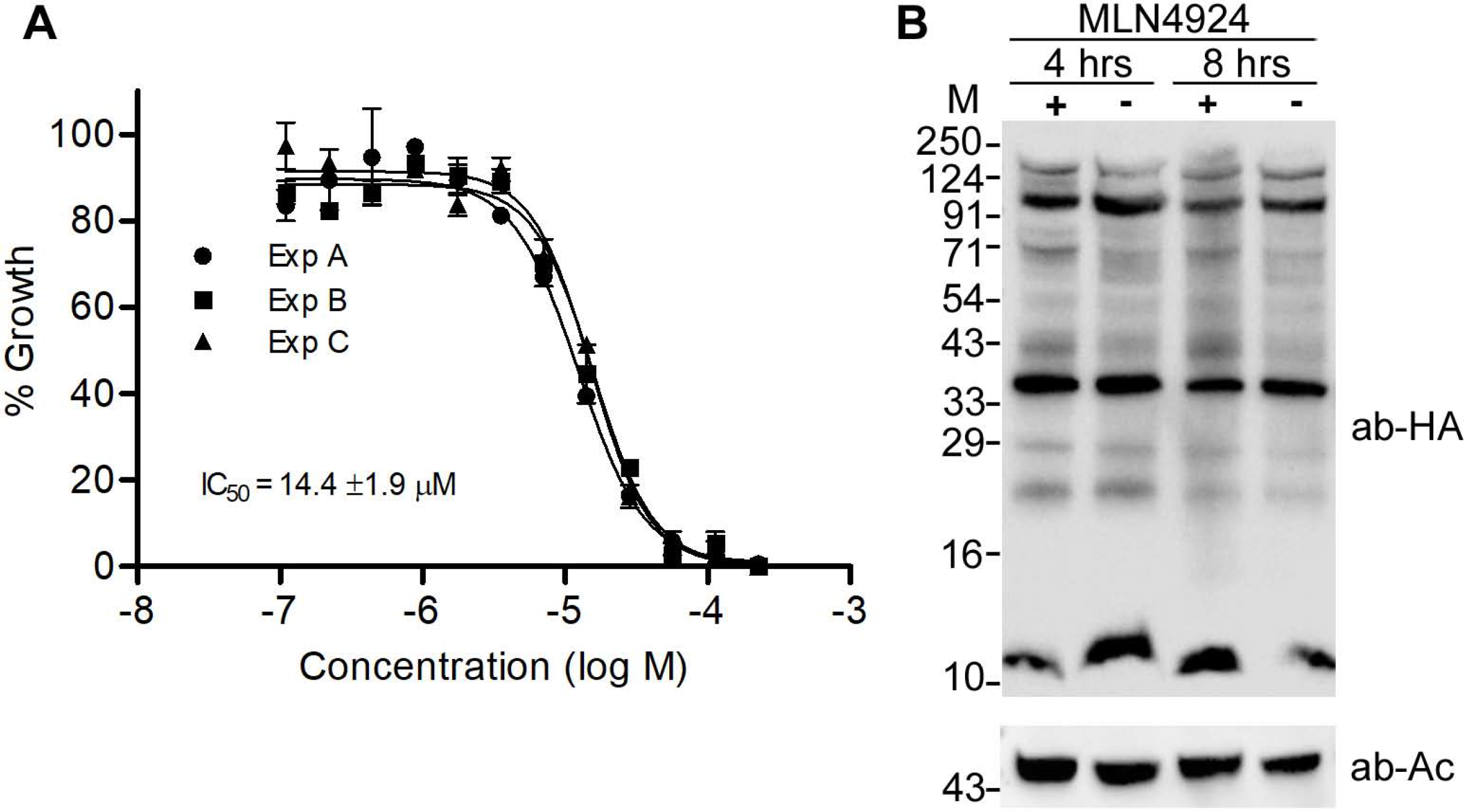
The effect of MLN4924 on neddylation. **A.** Wild type *P. falciparum* D10 ring stage parasites were incubated with various concentrations of MLN4924 for 48-50 hours, and the % growth (y-axis) at various MLN4924 concentrations was plotted to determine 50% inhibitory concentration (IC_50_) as described in the method section. The graph shows data from three independent experiments, with each point representing mean of two replicates. The IC_50_ value is average of three experiments with SD. **B.** The HA-PfNEDD8-expressing *P. falciparum* trophozoite stage parasites were grown with MLN4924 (+) or DMSO (-) for 4 or 8 hours, and then processed for western blotting using anti-HA antibodies (ab-HA). β-actin was used as a loading control (ab-Ac). The intensity of high molecular weight bands in MNL4924-treated parasites for both the time points appears to be comparable to that of DMSO-treated ones. The protein size markers are in kDa (M).

### PfNEDD8 substrates are *Plasmodium* cullins

Our in vitro neddylation assay and complementation experiments in *S. cerevisiae* demonstrated that cullins are the major substrates of PfNEDD8. To unequivocally prove that cullins are conjugated to PfNEDD8 in the parasite, we carried out immunoprecipitation of HN8-expressing and wild type parasite lysates using anti-HA antibodies and the immunoprecipitates were subjected to mass spectrometry. Two putative *P. falciparum* cullins (PF3D7_0811000 and PF3D7_0629800) were consistently identified in the HN8 immunoprecipitate, but not that of wild type parasites (Table 5, Fig. 7), confirming that *Plasmodium* cullins are biologically relevant substrates of PfNEDD8. The putative cullins are annotated as culin-1 (PF3D7_0811000) and cullin-like protein (PF3D7_0629800) in PlasmoDB. We have renamed the cullin-like protein as cullin-2. *P. falciparum* cullin-1 (Pfullin-1) is of the predicted size of 100 kDa and *P. falciparum* cullin-2 (Pfcullin-2) is of the predicated size of 137 kDa. The two prominent high molecular weight conjugated forms of PfNEDD8 are close to the neddylated Pfcullin-1 and Pfcullin-2 (Fig. 5A and Fig. 7). Apart from cullins, HN8 immunoprecipitate also contained ADP-ribosyl factor-1 (ARF1), which is an essential component of vesicular trafficking [97]. ARF1 does not seem to contain a putative neddylation motif and it is difficult to conclude at this point whether it is a direct NEDD8 substrate. It may be associated with some of the neddylation substrates.

**Table 5:** Proteins identified in the HN8 immunoprecipitate from parasites. HN8 and WT *P. falciparum* lysates were immunoprecipited and the eluates were subjected to mass spectrometry. The proteins listed are exclusively identified in the HN8 immunoprecipitate at least three of the 4 biological replicates. Shown above are the score (Sc), coverage (Co), and number of unique peptides (UP) for each protein. The predicted/reported role is based on their homologs in other systems.

| ID | Protein | Experiment 1 |  |  | Experiment 2 |  |  | Experiment 3 |  |  |
| --- | --- | --- | --- | --- | --- | --- | --- | --- | --- | --- |
|  |  | Sc | Co | UP | Sc | Co | UP | Sc | Co | UP |
| Q8IEI4<br>(PF3D7_1313000) | Ubiquitin-like protein nedd8 homologue | 3.53 | 19.74 | 1 | 2.62 | 35.53 | 1 | 3.27 | 51.32 | 1 |
| Q8IAU5<br>(PF3D7_0811000) | Cullin-1 | 13.92 | 8.44 | 7 | 14.32 | 21.95 | 9 | 14 | 22.8 | - |
| C6KTD3<br>(PF3D7_0629800) | Cullin-like protein | 15.42 | 7.53 | 7 | 4.67 | 10.89 | 3 | 2.91 | 20.11 | 4 |
| Q7KQL3<br>(PF3D7_1020900) | ADP-ribosylation factor 1 | 3.29 | 14.36 | 2 | 12.04 | 6.08 | 1 | 7.09 | 27.07 | - |

**Fig. 7.**
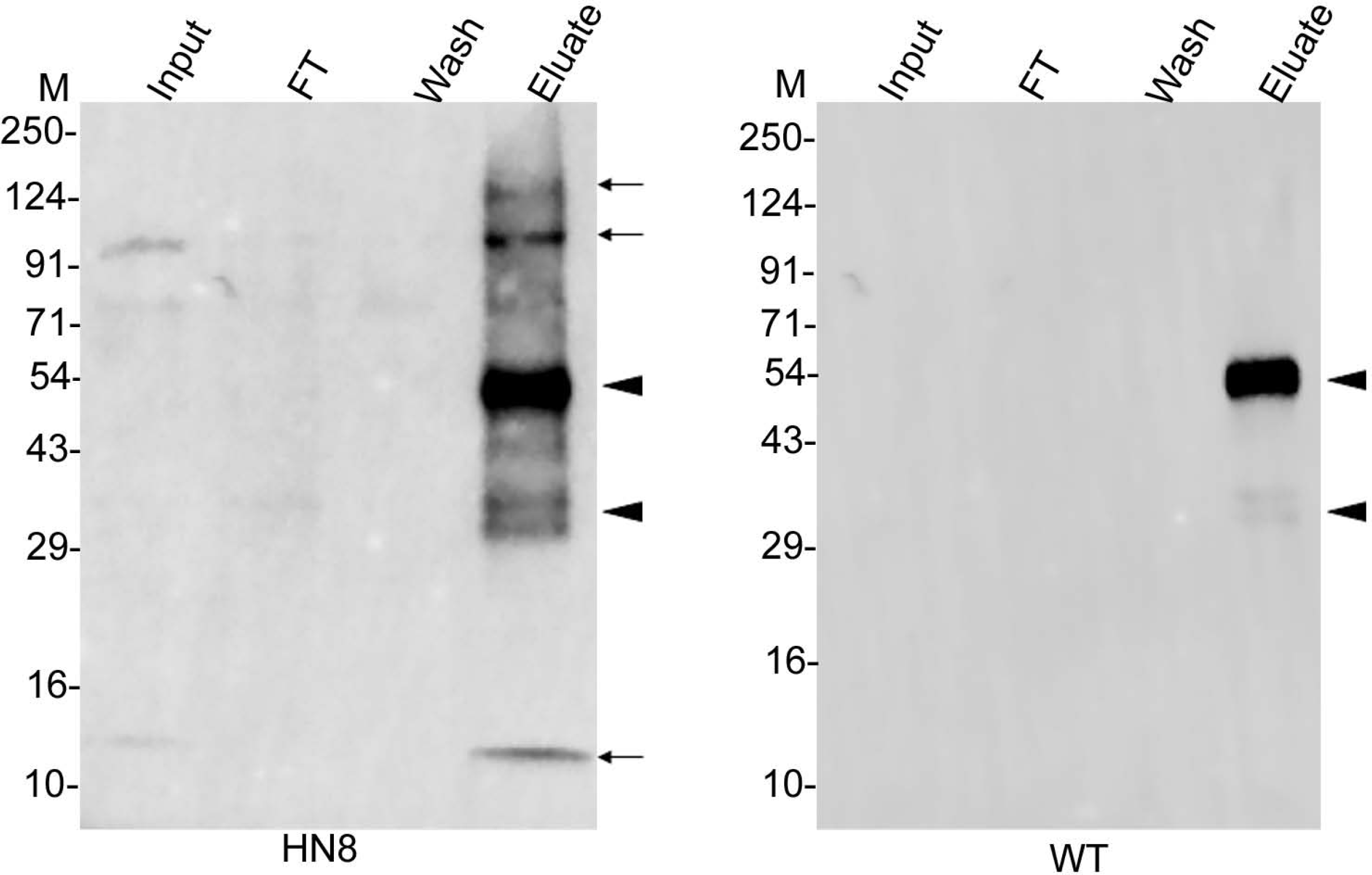
Western blot of parasite immunoprecipitates. The lysates of HA-PfNEDD8-expressing (HN8) and wild type (WT) parasites were immunoprecipitated with rabbit anti-HA antibodies. The eluates (Eluate) along with the extract (Input), flow through (FT) and wash (Wash) were analyzed by western blotting using mouse anti-HA antibodies. The HN8 blot shows the bands corresponding to free HA-PfNEDD8 and high molecular weight conjugates (arrow). The arrow heads indicate non-specific signal in both the blots. The protein size markers are in kDa (M).

In this study, we demonstrate for the first time that *P. falciparum* has a functional neddylation pathway. Given that neddylation is crucial for cell survival, particularly as a regulator of SCF E3 ubiquitin ligase that is critical for cell cycle, the characterization of *Plasmodium falciparum* NEDD8 and the identification of cullins as neddylation substrates lay down ground for investigation of specific roles and drug target potential of neddylation pathway in apicomplexan parasites.

## ACKNOWLEDGEMENTS

This study and the salaries of NA and DD were supported by funding from the Department of Biotechnology, India (grant reference numbers: 102/IFD/SAN/2914/2015-2016 and BT/PR11497/MED/29/854/2014) and the Council of Scientific and Industrial Research, India. M.B is a recipient of the Senior Research Fellowship from CSIR, India. RS and ZR were supported by fellowships from DBT. The authors would like to acknowledge CCMB central microscopy and proteomics facility for their technical support. The authors thank Dr. H.H. Krishnan for his valuable inputs as a Doctoral Advisory Committee member of MB.

## AUTHOR CONTRIBUTIONS

MB, NA and PSS conceived the project, designed experiments, interpreted the results and wrote the manuscript. MB and NA carried out the majority of experiments. RS, ZR and DD contributed to immunoprecipitation, mass spectrometry and cloning experiments. PMR guided NA for *S. cerevisiae* experiments.

## REFERENCES

1 Hochstrasser, M. (2009) Origin and function of ubiquitin-like proteins. Nature, Nature Publishing Group 458, 422–429.

2 Cappadocia, L., Lima, C. D. and Hochstrasser, M. (2018) Ubiquitin-like protein conjugation: structures, chemistry, and mechanism. Nature, ACS Publications 118, 889–918.

3 Cappadocia, L. and Lima, C. D. (2018) Ubiquitin-like protein conjugation: structures, chemistry, and mechanism. Chem. Rev., ACS Publications 118, 889–918.

4 Welchman, R. L., Gordon, C. and Mayer, R. J. (2005) Ubiquitin and ubiquitin-like proteins as multifunctional signals. Nat. Rev. Mol. cell Biol., Nature Publishing Group 6, 599–609.

5 Walsh, C. (2006) Posttranslational modification of proteins: expanding nature’s inventory, Roberts and Company Publishers.

6 Organization, W. H. (2019) World malaria report 2019, World Health Organization.

7 Menard, D. and Dondorp, A. (2017) Antimalarial drug resistance: a threat to malaria elimination. Cold Spring Harb. Perspect. Med., Cold Spring Harbor Laboratory Press 7, a025619.

8 Plowe, C. V, Cortese, J. F., Djimde, A., Nwanyanwu, O. C., Watkins, W. M., Winstanley, P. A., Franco, J. G. E., Mollinedo, R. E., Avila, J. C. and Cespedes, J. L. (1997) Mutations in Plasmodium falciparum dihydrofolate reductase and dihydropteroate synthase and epidemiologic patterns of pyrimethamine-sulfadoxine use and resistance. J. Infect. Dis., The University of Chicago Press 176, 1590–1596.

9 Dondorp, A. M., Nosten, F., Yi, P., Das, D., Phyo, A. P., Tarning, J., Lwin, K. M., Ariey, F., Hanpithakpong, W. and Lee, S. J. (2009) Artemisinin resistance in Plasmodium falciparum malaria. N. Engl. J. Med., Mass Medical Soc 361, 455–467.

10 Korsinczky, M., Chen, N., Kotecka, B., Saul, A., Rieckmann, K. and Cheng, Q. (2000) Mutations in Plasmodium falciparumCytochrome b That Are Associated with Atovaquone Resistance Are Located at a Putative Drug-Binding Site. Antimicrob. Agents Chemother., Am Soc Microbiol 44, 2100–2108.

11 Fidock, D. A., Nomura, T., Talley, A. K., Cooper, R. A., Dzekunov, S. M., Ferdig, M. T., Ursos, L. M. B., Naudé, B., Deitsch, K. W. and Su, X. (2000) Mutations in the P. falciparum digestive vacuole transmembrane protein PfCRT and evidence for their role in chloroquine resistance. Mol. Cell, Elsevier 6, 861–871.

12 Ponder, E. L. and Bogyo, M. (2007) Ubiquitin-like modifiers and their deconjugating enzymes in medically important parasitic protozoa. Eukaryot. Cell, Am Soc Microbiol 6, 1943–1952.

13 Hall, N., Pain, A., Berriman, M., Churcher, C., Harris, B., Harris, D., Mungall, K., Bowman, S., Atkin, R. and Baker, S. (2002) Sequence of Plasmodium falciparum chromosomes 1, 3–9 and 13. Nature, Nature Publishing Group 419, 527–531.

14 Cosson, P., Demolliere, C., Hennecke, S., Duden, R., Letourneur, F., Korsinczky, M., Chen, N., Kotecka, B., Saul, A., Rieckmann, K., et al. (2009) Deciphering the ubiquitin-mediated pathway in apicomplexan parasites: a potential strategy to interfere with parasite virulence. Cell, Elsevier, United States 176, 861–871.

15 Prasad, R., Atul, V. K. K., Legac, J., Singhal, N., Navale, R., Rosenthal, P. J. and Sijwali, P. S. (2013) Blocking Plasmodium falciparum development via dual inhibition of hemoglobin degradation and the ubiquitin proteasome system by MG132. PLoS One, Public Library of Science 8.

16 Ng, C. L., Fidock, D. A. and Bogyo, M. (2017) Protein degradation systems as antimalarial therapeutic targets. Trends Parasitol., Elsevier 33, 731–743.

17 Dogovski, C., Xie, S. C., Burgio, G., Bridgford, J., Mok, S., McCaw, J. M., Chotivanich, K., Kenny, S., Gnädig, N. and Straimer, J. (2015) Targeting the cell stress response of Plasmodium falciparum to overcome artemisinin resistance. PLoS Biol., Public Library of Science 13.

18 Reiter, K., Mukhopadhyay, D., Zhang, H., Boucher, L. E., Kumar, N., Bosch, J. and Matunis, M. J. (2013) Identification of biochemically distinct properties of the small ubiquitin-related modifier (SUMO) conjugation pathway in Plasmodium falciparum. J. Biol. Chem., ASBMB 288, 27724–27736.

19 Navale, R., Atul, A. D. A. and Sijwali, P. S. (2014) Characterization of the autophagy marker protein Atg8 reveals atypical features of autophagy in Plasmodium falciparum. PLoS One, Public Library of Science 9.

20 Walczak, M., Ganesan, S. M., Niles, J. C. and Yeh, E. (2018) ATG8 is essential specifically for an autophagy-independent function in apicoplast biogenesis in bloodstage malaria parasites. MBio, Am Soc Microbiol 9.

21 Kitamura, K., Kishi-Itakura, C., Tsuboi, T., Sato, S., Kita, K., Ohta, N. and Mizushima, N. (2012) Autophagy-related Atg8 localizes to the apicoplast of the human malaria parasite Plasmodium falciparum. PLoS One, Public Library of Science 7, e42977.

22 Tomlins, A. M., Ben-Rached, F., Williams, R. A. M., Proto, W. R., Coppens, I., Ruch, U., Gilberger, T. W., Coombs, G. H., Mottram, J. C. and Müller, S. (2013) Plasmodium falciparum ATG8 implicated in both autophagy and apicoplast formation. Autophagy, Taylor & Francis 9, 1540–1552.

23 Cervantes, S., Bunnik, E. M., Saraf, A., Conner, C. M., Escalante, A., Sardiu, M. E., Ponts, N., Prudhomme, J., Florens, L. and Le Roch, K. G. (2014) The multifunctional autophagy pathway in the human malaria parasite, Plasmodium falciparum. Autophagy, Taylor & Francis 10, 80–92.

24 Wada, H., Kito, K., Caskey, L. S., Yeh, E. T. H. and Kamitani, T. (1998) Cleavage of the C-terminus of NEDD8 by UCH-L3. Biochem. Biophys. Res. Commun., Elsevier 251, 688–692.

25 Shen, L., Liu, H., Dong, C., Xirodimas, D., Naismith, J. H. and Hay, R. T. (2005) Structural basis of NEDD8 ubiquitin discrimination by the deNEDDylating enzyme NEDP1. EMBO J., John Wiley & Sons, Ltd 24, 1341–1351.

26 Walden, H., Podgorski, M. S., Huang, D. T., Miller, D. W., Howard, R. J., Minor, D. L. J., Holton, J. M. and Schulman, B. A. (2003) The structure of the APPBP1-UBA3-NEDD8-ATP complex reveals the basis for selective ubiquitin-like protein activation by an E1. Mol. Cell, United States 12, 1427–1437.

27 Gong, L. and Yeh, E. T. H. (1999) Identification of the activating and conjugating enzymes of the NEDD8 conjugation pathway. J. Biol. Chem., ASBMB 274, 12036–12042.

28 Kamura, T., Conrad, M. N., Yan, Q., Conaway, R. C. and Conaway, J. W. (1999) The Rbx1 subunit of SCF and VHL E3 ubiquitin ligase activates Rub1 modification of cullins Cdc53 and Cul2. Genes Dev., Cold Spring Harbor Lab 13, 2928–2933.

29 Ma, T., Chen, Y., Zhang, F., Yang, C.-Y., Wang, S. and Yu, X. (2013) RNF111-dependent neddylation activates DNA damage-induced ubiquitination. Mol. Cell, Elsevier 49, 897–907.

30 Kurz, T., Chou, Y.-C., Willems, A. R., Meyer-Schaller, N., Hecht, M.-L., Tyers, M., Peter, M. and Sicheri, F. (2008) Dcn1 functions as a scaffold-type E3 ligase for cullin neddylation. Mol. Cell, Elsevier 29, 23–35.

31 Xirodimas, D. P., Saville, M. K., Bourdon, J.-C., Hay, R. T. and Lane, D. P. (2004) Mdm2-mediated NEDD8 conjugation of p53 inhibits its transcriptional activity. Cell, Elsevier 118, 83–97.

32 Santonico, E. (2019) New insights into the mechanisms underlying NEDD8 structural and functional specificities. In Ubiquitin Proteasome System-Current Insights into Mechanism Cellular Regulation and Disease, IntechOpen.

33 Dharmasiri, S., Dharmasiri, N., Hellmann, H. and Estelle, M. (2003) The RUB/Nedd8 conjugation pathway is required for early development in Arabidopsis. EMBO J., John Wiley & Sons, Ltd Chichester, UK 22, 1762–1770.

34 Osaka, F., Saeki, M., Katayama, S., Aida, N., Toh-e, A., Kominami, K., Toda, T., Suzuki, T., Chiba, T. and Tanaka, K. (2000) Covalent modifier NEDD8 is essential for SCF ubiquitin-ligase in fission yeast. EMBO J., John Wiley & Sons, Ltd 19, 3475–3484.

35 Jones, D. and Candido, E. P. M. (2000) The NED-8 conjugating system in Caenorhabditis elegans is required for embryogenesis and terminal differentiation of the hypodermis. Dev. Biol., Elsevier 226, 152–165.

36 Tateishi, K., Omata, M., Tanaka, K. and Chiba, T. (2001) The NEDD8 system is essential for cell cycle progression and morphogenetic pathway in mice. J. Cell Biol., Rockefeller University Press 155, 571–580.

37 Ou, C.-Y., Lin, Y.-F., Chen, Y.-J. and Chien, C.-T. (2002) Distinct protein degradation mechanisms mediated by Cul1 and Cul3 controlling Ci stability in Drosophila eye development. Genes Dev., Cold Spring Harbor Lab 16, 2403–2414.

38 Yu, G., Liu, X., Zhang, D., Wang, J., Ouyang, G., Chen, Z. and Xiao, W. (2019) Zebrafish Nedd8 Facilitates Ovarian Development and the Maintenance of Female Secondary Sexual Characteristics Via Suppression of Androgen Receptor Activity, Facilitates Ovarian Development and the Maintenance of Female Secondary ….

39 Hori, T., Osaka, F., Chiba, T., Miyamoto, C., Okabayashi, K., Shimbara, N., Kato, S. and Tanaka, K. (1999) Covalent modification of all members of human cullin family proteins by NEDD8. Oncogene, Nature Publishing Group 18, 6829–6834.

40 Lammer, D., Mathias, N., Laplaza, J. M., Jiang, W., Liu, Y., Callis, J., Goebl, M. and Estelle, M. (1998) Modification of yeast Cdc53p by the ubiquitin-related protein rub1p affects function of the SCFCdc4 complex. Genes Dev., Cold Spring Harbor Lab 12, 914–926.

41 Carrabino, S., Carminati, E., Talarico, D., Pardi, R. and Bianchi, E. (2004) Expression pattern of the JAB1/CSN5 gene during murine embryogenesis: colocalization with NEDD8. Gene Expr. patterns, Elsevier 4, 423–431.

42 Gao, F., Cheng, J., Shi, T. and Yeh, E. T. H. (2006) Neddylation of a breast cancer-associated protein recruits a class III histone deacetylase that represses NFκB-dependent transcription. Nat. Cell Biol., Nature Publishing Group 8, 1171–1177.

43 Liu, J., Furukawa, M., Matsumoto, T. and Xiong, Y. (2002) NEDD8 modification of CUL1 dissociates p120CAND1, an inhibitor of CUL1-SKP1 binding and SCF ligases. Mol. Cell, Elsevier 10, 1511–1518.

44 Bosu, D. R. and Kipreos, E. T. (2008) Cullin-RING ubiquitin ligases: global regulation and activation cycles. Cell Div., BioMed Central 3, 7.

45 Schwechheimer, C. and Mergner, J. (2014) The NEDD8 modification pathway in plants. Front. Plant Sci. 5, 103.

46 Sufan, R. I. and Ohh, M. (2006) Role of the NEDD8 modification of Cul2 in the sequential activation of ECV complex. Neoplasia (New York, NY), Neoplasia Press 8, 956.

47 Brown, J. S., Lukashchuk, N., Sczaniecka-Clift, M., Britton, S., le Sage, C., Calsou, P., Beli, P., Galanty, Y. and Jackson, S. P. (2015) Neddylation promotes ubiquitylation and release of Ku from DNA-damage sites. Cell Rep., Elsevier 11, 704–714.

48 Duda, D. M., Borg, L. A., Scott, D. C., Hunt, H. W., Hammel, M. and Schulman, B. A. (2008) Structural insights into NEDD8 activation of cullin-RING ligases: conformational control of conjugation. Cell, Elsevier 134, 995–1006.

49 Merlet, J., Burger, J., Gomes, J.-E. and Pintard, L. (2009) Regulation of cullin-RING E3 ubiquitin-ligases by neddylation and dimerization. Cell. Mol. life Sci., Springer 66, 1924–1938.

50 Morimoto, M., Nishida, T., Honda, R. and Yasuda, H. (2000) Modification of cullin-1 by ubiquitin-like protein Nedd8 enhances the activity of SCFskp2 toward p27kip1. Biochem. Biophys. Res. Commun., Academic Press 270, 1093–1096.

51 Oved, S., Mosesson, Y., Zwang, Y., Santonico, E., Shtiegman, K., Marmor, M. D., Kochupurakkal, B. S., Katz, M., Lavi, S. and Cesareni, G. (2006) Conjugation to Nedd8 instigates ubiquitylation and down-regulation of activated receptor tyrosine kinases. J. Biol. Chem., ASBMB 281, 21640–21651.

52 Zhou, L., Zhang, W., Sun, Y. and Jia, L. (2018) Protein neddylation and its alterations in human cancers for targeted therapy. Cell. Signal., Elsevier 44, 92–102.

53 Barbier-Torres, L., Delgado, T. C., García-Rodríguez, J. L., Zubiete-Franco, I., Fernández-Ramos, D., Buqué, X., Cano, A., Gutiérrez-de Juan, V., Fernández-Domínguez, I. and Lopitz-Otsoa, F. (2015) Stabilization of LKB1 and Akt by neddylation regulates energy metabolism in liver cancer. Oncotarget, Impact Journals, LLC 6, 2509.

54 Wang, M., Medeiros, B. C., Erba, H. P., DeAngelo, D. J., Giles, F. J. and Swords, R. T. (2011) Targeting protein neddylation: a novel therapeutic strategy for the treatment of cancer. Expert Opin. Ther. Targets, Taylor & Francis 15, 253–264.

55 Swords, R. T., Erba, H. P., DeAngelo, D. J., Bixby, D. L., Altman, J. K., Maris, M., Hua, Z., Blakemore, S. J., Faessel, H. and Sedarati, F. (2015) Pevonedistat (MLN 4924), a First-in-Class NEDD 8-activating enzyme inhibitor, in patients with acute myeloid leukaemia and myelodysplastic syndromes: a phase 1 study. Br. J. Haematol., Wiley Online Library 169, 534–543.

56 Nawrocki, S. T., Kelly, K. R., Smith, P. G., Espitia, C. M., Possemato, A., Beausoleil, S. A., Milhollen, M., Blakemore, S., Thomas, M. and Berger, A. (2013) Disrupting protein NEDDylation with MLN4924 is a novel strategy to target cisplatin resistance in ovarian cancer. Clin. cancer Res., AACR 19, 3577–3590.

57 Liao, S., Hu, H., Wang, T., Tu, X. and Li, Z. (2017) The Protein Neddylation Pathway in Trypanosoma brucei FUNCTIONAL CHARACTERIZATION AND SUBSTRATE IDENTIFICATION. J. Biol. Chem., ASBMB 292, 1081–1091.

58 Artavanis-Tsakonas, K., Misaghi, S., Comeaux, C. A., Catic, A., Spooner, E., Duraisingh, M. T. and Ploegh, H. L. (2006) Identification by functional proteomics of a deubiquitinating/deNeddylating enzyme in Plasmodium falciparum. Mol. Microbiol., Wiley Online Library 61, 1187–1195.

59 Frickel, E., Quesada, V., Muething, L., Gubbels, M., Spooner, E., Ploegh, H. and Artavanis-Tsakonas, K. (2007) Apicomplexan UCHL3 retains dual specificity for ubiquitin and Nedd8 throughout evolution. Cell. Microbiol., Wiley Online Library 9, 1601–1610.

60 Artavanis-Tsakonas, K., Weihofen, W. A., Antos, J. M., Coleman, B. I., Comeaux, C. A., Duraisingh, M. T., Gaudet, R. and Ploegh, H. L. (2010) Characterization and structural studies of the Plasmodium falciparum ubiquitin and Nedd8 hydrolase UCHL3. J. Biol. Chem., ASBMB 285, 6857–6866.

61 Karpiyevich, M., Adjalley, S., Mol, M., Ascher, D. B., Mason, B., van der Heden van Noort, G. J., Laman, H., Ovaa, H., Lee, M. C. S. and Artavanis-Tsakonas, K. (2019) Nedd8 hydrolysis by UCH proteases in Plasmodium parasites. PLOS Pathog., Public Library of Science 15, e1008086.

62 Altschul, S. F., Gish, W., Miller, W., Myers, E. W. and Lipman, D. J. (1990) Basic local alignment search tool. J. Mol. Biol., Elsevier 215, 403–410.

63 Aurrecoechea, C., Brestelli, J., Brunk, B. P., Dommer, J., Fischer, S., Gajria, B., Gao, X., Gingle, A., Grant, G., Harb, O. S., et al. (2009) PlasmoDB: a functional genomic database for malaria parasites. Nucleic Acids Res. 37, D539–D543.

64 Sievers, F., Wilm, A., Dineen, D., Gibson, T. J., Karplus, K., Li, W., Lopez, R., McWilliam, H., Remmert, M. and Söding, J. (2011) Fast, scalable generation of high-quality protein multiple sequence alignments using Clustal Omega. Mol. Syst. Biol., John Wiley & Sons, Ltd 7.

65 Armougom, F., Moretti, S., Poirot, O., Audic, S., Dumas, P., Schaeli, B., Keduas, V. and Notredame, C. (2006) Expresso: automatic incorporation of structural information in multiple sequence alignments using 3D-Coffee. Nucleic Acids Res., Oxford University Press 34, W604–W608.

66 Kumar, S., Stecher, G., Li, M., Knyaz, C. and Tamura, K. (2018) MEGA X: molecular evolutionary genetics analysis across computing platforms. Mol. Biol. Evol., Oxford University Press 35, 1547–1549.

67 Jones, D. T., Taylor, W. R. and Thornton, J. M. (1992) The rapid generation of mutation data matrices from protein sequences. Bioinformatics, Oxford University Press 8, 275–282.

68 Trager, W. and Jensen, J. B. (1976) Human malaria parasites in continuous culture. Science (80-.)., American Association for the Advancement of Science 193, 673–675.

69 Lambros, C. and Vanderberg, J. (1979) Synchronization of Plasmodium Falciparum Erythrocytic Stages in Culture - PubMed. J. Parasitol. 65, 418–420.

70 Nair, D. N., Prasad, R., Singhal, N., Bhattacharjee, M., Sudhakar, R., Singh, P., Thanumalayan, S., Kiran, U., Sharma, Y. and Sijwali, P. S. (2018) A conserved human DJ1-subfamily motif (DJSM) is critical for anti-oxidative and deglycase activities of Plasmodium falciparum DJ1. Mol. Biochem. Parasitol., Elsevier 222, 70–80.

71 Fidock, D. A. and Wellems, T. E. (1997) Transformation with human dihydrofolate reductase renders malaria parasites insensitive to WR99210 but does not affect the intrinsic activity of proguanil. Proc. Natl. Acad. Sci., National Acad Sciences 94, 10931–10936.

72 Sijwali, P. S. and Rosenthal, P. J. (2004) Gene disruption confirms a critical role for the cysteine protease falcipain-2 in hemoglobin hydrolysis by Plasmodium falciparum. Proc. Natl. Acad. Sci. U. S. A., National Academy of Sciences 101, 4384–4389.

73 Govindarajalu, G., Rizvi, Z., Kumar, D. and Sijwali, P. S. (2019) Lyse-Reseal erythrocytes for transfection of Plasmodium falciparum. Sci. Rep., Nature Publishing Group 9, 1–8.

74 Wu, Y., Sifri, C. D., Lei, H. H., Su, X. Z. and Wellems, T. E. (1995) Transfection of Plasmodium falciparum within human red blood cells. Proc. Natl. Acad. Sci. U. S. A. 92, 973–977.

75 Van den Hoff, M. J., Moorman, A. F. and Lamers, W. H. (1992) Electroporation in’intracellular’buffer increases cell survival. Nucleic Acids Res., Oxford University Press 20, 2902.

76 Chiba, T. (2005) In vitro systems for NEDD8 conjugation by Ubc12. Methods Enzymol., United States 398, 68–73.

77 Leidecker, O. and Xirodimas, D. P. (2012) Isolation of NEDDylated proteins in human cells. Methods Mol. Biol., United States 832, 133–140.

78 Janke, C., Magiera, M. M., Rathfelder, N., Taxis, C., Reber, S., Maekawa, H., Moreno-Borchart, A., Doenges, G., Schwob, E., Schiebel, E., et al. (2004) A versatile toolbox for PCR-based tagging of yeast genes: New fluorescent proteins, more markers and promoter substitution cassettes. Yeast 21, 947–962.

79 Knop, M., Siegers, K., Pereira, G., Zachariae, W., Winsor, B., Nasmyth, K. and Schiebel, E. (1999) Epitope tagging of yeast genes using a PCR-based strategy: More tags and improved practical routines. Yeast 15, 963–972.

80 Hoffman, C. S. and Winston, F. (1987) A ten-minute DNA preparation from yeast efficiently releases autonomous plasmids for transformaion of Escherichia coli. Gene 57, 267–272.

81 Kandasamy, G. and Andréasson, C. (2018) Hsp70-Hsp110 chaperones deliver ubiquitin-dependent and - independent substrates to the 26S proteasome for proteolysis in yeast. J. Cell Sci., Company of Biologists Ltd 131.

82 Whitby, F. G., Xia, G., Pickart, C. M. and Hill, C. P. (1998) Crystal structure of the human ubiquitin-like protein NEDD8 and interactions with ubiquitin pathway enzymes. J. Biol. Chem., ASBMB 273, 34983–34991.

83 Huang, D. T., Hunt, H. W., Zhuang, M., Ohi, M. D., Holton, J. M. and Schulman, B. A. (2007) Basis for a ubiquitin-like protein thioester switch toggling E1–E2 affinity. Nature, Nature Publishing Group 445, 394–398.

84 Souphron, J., Waddell, M. B., Paydar, A., Tokgöz-Gromley, Z., Roussel, M. F. and Schulman, B. A. (2008) Structural dissection of a gating mechanism preventing misactivation of ubiquitin by NEDD8’s E1. Biochemistry, ACS Publications 47, 8961–8969.

85 Ciechanover, Aeshh., Elias, S., Heller, H. and Hershko, A. (1982) “Covalent affinity” purification of ubiquitin-activating enzyme. J. Biol. Chem., ASBMB 257, 2537–2542.

86 Schulman, B. A. and Harper, J. W. (2009) Ubiquitin-like protein activation by E1 enzymes: the apex for downstream signalling pathways. Nat. Rev. Mol. cell Biol., Nature Publishing Group 10, 319–331.

87 Bai, C., Sen, P., Hofmann, K., Ma, L., Goebl, M., Harper, J. W. and Elledge, S. J. (1996) SKP1 connects cell cycle regulators to the ubiquitin proteolysis machinery through a novel motif, the F-box. Cell, Elsevier 86, 263–274.

88 Zheng, N., Schulman, B. A., Song, L., Miller, J. J., Jeffrey, P. D., Wang, P., Chu, C., Koepp, D. M., Elledge, S. J. and Pagano, M. (2002) Structure of the Cul1–Rbx1– Skp1–F box Skp2 SCF ubiquitin ligase complex. Nature, Nature Publishing Group 416, 703–709.

89 Liakopoulos, D., Doenges, G., Matuschewski, K. and Jentsch, S. (1998) A novel protein modification pathway related to the ubiquitin system. EMBO J., John Wiley & Sons, Ltd 17, 2208–2214.

90 Soucy, T. A., Smith, P. G., Milhollen, M. A., Berger, A. J., Gavin, J. M., Adhikari, S., Brownell, J. E., Burke, K. E., Cardin, D. P., Critchley, S., et al. (2009) An inhibitor of NEDD8-activating enzyme as a new approach to treat cancer. Nature, England 458, 732–736.

91 Brownell, J. E., Sintchak, M. D., Gavin, J. M., Liao, H., Bruzzese, F. J., Bump, N. J., Soucy, T. A., Milhollen, M. A., Yang, X. and Burkhardt, A. L. (2010) Substrate-assisted inhibition of ubiquitin-like protein-activating enzymes: the NEDD8 E1 inhibitor MLN4924 forms a NEDD8-AMP mimetic in situ. Mol. Cell, Elsevier 37, 102–111.

92 Soucy, T. A., Dick, L. R., Smith, P. G., Milhollen, M. A. and Brownell, J. E. (2010) The NEDD8 Conjugation Pathway and Its Relevance in Cancer Biology and Therapy. Genes Cancer, SAGE Publications 1, 708–716.

93 Milhollen, M. A., Traore, T., Adams-Duffy, J., Thomas, M. P., Berger, A. J., Dang, L., Dick, L. R., Garnsey, J. J., Koenig, E. and Langston, S. P. (2010) MLN4924, a NEDD8-activating enzyme inhibitor, is active in diffuse large B-cell lymphoma models: rationale for treatment of NF-κB–dependent lymphoma. Blood, J. Am. Soc. Hematol., American Society of Hematology Washington, DC 116, 1515–1523.

94 Aubry, A., Yu, T. and Bremner, R. (2020) Preclinical studies reveal MLN4924 is a promising new retinoblastoma therapy. Cell Death Discov., Nature Publishing Group 6, 1–12.

95 Lin, S., Shang, Z., Li, S., Gao, P., Zhang, Y., Hou, S., Qin, P., Dong, Z., Hu, T. and Chen, P. (2018) Neddylation inhibitor MLN4924 induces G(2) cell cycle arrest, DNA damage and sensitizes esophageal squamous cell carcinoma cells to cisplatin. Oncol. Lett., D.A. Spandidos 15, 2583–2589.

96 Yoshimura, C., Muraoka, H., Ochiiwa, H., Tsuji, S., Hashimoto, A., Kazuno, H., Nakagawa, F., Komiya, Y., Suzuki, S. and Takenaka, T. (2019) TAS4464, a highly potent and selective inhibitor of NEDD8-activating enzyme, suppresses neddylation and shows antitumor activity in diverse cancer models. Mol. Cancer Ther., AACR 18, 1205–1216.

97 Cook, W. J., Smith, C. D., Senkovich, O., Holder, A. A. and Chattopadhyay, D. (2010) Structure of Plasmodium falciparum ADP-ribosylation factor 1. Acta Crystallogr. Sect. F, Wiley Online Library 66, 1426–1431.

