## Supplementary information for "Characterization of *Plasmodium falciparum* NEDD8 and identification of cullins as its substrates"

**Table S1**. List of the primers used in the study. The sequences are in 5'-3' direction and restriction enzyme sites are in bold.

| | |
|---|---|
| PfHN8-F | AATT**AGATCT**CAAAATGTATCCATATGATGTTCCAGATTACGCTGGAAGTCAAATATTAGTTAAAACATTAACAGGG (HA-tag is underlined) |
| PfN8ex-R | ATT**CTCGAG**TTATCCTCCTCTTAATTG |
| PfN8ggM-R | AAT**CTCGAG**TTATGCTGCTCTTAATTG (codons for Gly75-Gly76 substitution to Ala-Ala are underlined) |
| SN8-F | AGAGAACAGATTGGAGGCATGCAAATATTAGTTAAAACATTAACAG |
| SN8-R | CTCGAGTGCGGCCGC**AAGCTT**TTATCCTCCTCTTAATTGTAAAATT |
| SMT3-F | ATTA**CATATG**CATCACCATCACCATCACTCGGACTCAGAAGTCAATCAA (His-tag sequence is underlined) |
| SMT3-R | AATC**AAGCTT**AAATACGTAGGCCTCCAATCTGTTCTCTGTGAGC |
| ScN8KO-F | CGACAGAGGAATAAATAAAGGAAGGTAATTAACTTCCTTACAGCCGTAACCGATGCGTACGCTGCAGGTCGAC (sequence of ScRub1 locus is underlined) |
| ScN8KO-R | TCTTTTCTAATGAACACCTTCGATAAAATTCCATAAATGACGGAAAATGGTTCTAATCGATGAATTCGAGCTCG (sequence of ScRub1 locus is underlined) |
| ScN8-Fcon | CTTCGGGAAGGCAAACGAACAAC |
| ScN8-Rcon | GTAGCCTTCCAAAGTCCAAGTGAAC |
| ScU12KO-F | TACTTAGAGACATTAAAGAATCAGTTAGAGAATATAAAACAAGATAATAAAAATGCGTACGCTGCAGGTCGAC (sequence of ScUbc12 locus is underlined) |
| ScU12KO-R | ATTTTATTGTTTATATATAATCAAATCAACCGAAAATTACTCTGAACTTGACTCAATCGATGAATTCGAGCTCG (sequence of ScUbc12 locus is underlined) |
| ScU12-Fcon | CAATCCTCTGGTAAATCTCATCT |
| ScU12-Rcon | ACTAACAGAAGATGGCGGTAGC |
| ScN8-Fepi | ACCGCCG**GAATTC**ATGATTGTTAAAGTGAAGACACTG |
| ScN8-Repi | ATGCGC**GGATCC**CTAGTTACCACCTCTTAGTGTTAATAC |
| PfN8-FScepi | AAAACG**GAATTC**ATGCAAATATTAGTTAAAACATTAACAG |
| PfN8-Rsp | TTAA**GGATCC**TTATCCTCCTCTTAATTGTAAAATC |

**Fig. S1. Sequence alignment and phylogenetic analysis of NEDD8 proteins.** The NEDD8 protein sequences of *Plasmodium* species (grey shaded) were compared with 139 NEDD8 protein sequences of indicated organisms representing metazoans, plants, fungi and protozoans. The ID of each NEDD8 protein is in the bracket. The conserved amino acid residues are in red font. The amino acid residues conserved or shared by at least 75% of the proteins are in red and green font, respectively. Also shown is the consensus sequence shared by at least 75% of the proteins, with hyphens representing variation at these positions. The C-terminal tail processing site in marked with an arrow and amino acid residues marked with asterisk have been demonstrated to confer specificity for neddylation enzymes. Amino acid residues in blue font are conserved across Plasmodium PfNEDD8 proteins and drastically differ from the consensus in physicochemical properties.

**Fig. S2. Expression of purification of recombinant PfNEDD8**. PfNEDD8 was expressed as a C-terminal fusion of His-Sumo in *E. coli*, the His-Sumo-PfNEDD8 fusion protein (SN8) was purified by Ni-NTA chromatography under native conditions. The Coomassie stained SDS-PAGE shows lysates of uninduced (Un) and induced (In) bacteria, soluble (Sol) and insoluble (Ins) fractions of the induced bacteria, and Ni-NTA purified SN8 (Pur). The protein size markers are in kDa (M).

**Figure S3: Growth comparison of WT, *rub1*Δ and *ubc12*Δ *S. cerevisiae* strains.** The *rub1*Δ, *ubc12*Δ and wild type (WT) *S. cerevisiae* strains were grown in YPD medium for 14 hours beginning with 0.1 OD. Aliquots were collected at every two hours for the measurement of cell density at $OD_{600}$. The graph shows growth as $OD_{600}$ on y-axis versus time (hours), and indicates comparable growth of all three strains.

```
                                                  10        20        30        40        50        60        70        80        90
                                                   |         |         |         |         |         |         |         |         |
C.depauperatus(A0A1E3JCZ1)          ---MIVKVKTLTGKEVDI-DVQPDMTIGKVKEKVEEKAGIPPVQQRLISGGKAMNDEKTIQDYKIKAAD--VIHLVLALRGGRA-------------
K.bestiolae(A0A1B9GAI6)             ---MIVKVKTLTGKEVDI-DVQPDMTINKVKERVEEKAGIPPVQQRLIFGGKAMADDKAISDYKINAGA--VIHLVLALRGGR--------------
K.mangroviensis(A0A1B9IIT2)         ---MIVKVKTLTGKEVDI-DVQPDMTINKVKERVEEKAGIPPVQQRLIFGGKAMADDKAIQDYKINAGA--VIHLVLALRGGR--------------
C.gattii(A0A095CBA7)                ---MIVKVKTLTGKEVDI-DVQPDMTISKVKERVEEKAGIPPVQQRLIFGGKAMGDDKTIQDYKIQAGA--AIHLVLALRGGRA-------------
C.neoformans(J9VQ81)                ---MIVKVKTLTGKEVDI-DVQPDMTISKVKERVEEKAGIPPVQQRLIFGGKAMGDDKTIQDYKIQAGA--AIHLVLALRGGRA-------------
T.mesenterica(A0A4V1M4Z3)           ---MLIKVKTLTGKEIEL-DVQPETKVSKVKENVEEKAGIPPIQQRLIFGGKAMNDEKELGFYGVPAGG--VIHLVLALRGGR--------------
L.edodes(A0A1Q3EGM6)                ---MLIKVRTLTGKELEI-DVDSDAKIFTIKEKVEEQQGIPPVQQRLIFGGKQLDDDKHITETNIVAGS--TLHLVLALRGGR--------------
A.parvum(A0A023FVX4)                ---MLIKVKTLTGKEIEI-DIEPTDKVERIKERVEEKEGIPPAQQRLIFSGKQMNDDKTAADYKVTGGS--VLHLVLALRGG-EWCACYL-------
A.americanum(B5M745)                ---MLIKVKTLTGKEIEI-DIEPTDKVERIKERVEEKEGIPPAQQRLIFSGKQMNDDKTAADYKVTGGS--VLHLVLALRGG-EWRACNQ-------
H.vulgaris(T2MDB3)                  ---MLIKVKTLTGKEIEI-DIEPNDKVERIKERVEEKEGIPPAQQRLIFSGKQMNDDKTAQDYKVSGGS--VLHLVLALRGG---------------
A.ventricosus(A0A4Y2GFW1)           ---MLIKVKTLTGKEIEI-DIEPTDKVERIKERVEEKEGIPPAQQRLIFSGKQMNDDKTAFDYKVTGGS--VLHLVLALRGGYPSESCFC-------
A.cerana(A0A2A3E9N1)                ---MLIKVKTLTGKEIEI-DIEPTDKVERIKERVEEKEGIPPQQQRLIFSGKQMNDEKTAQDYKVQGGS--VLHLVLALRGGL--------------
A.planipennis(A0A1W4WES4)           ---MLIKVKTLTGKEIEI-DIEPTDKVERIKERVEEKEGIPPQQQRLIFSGKQMNDEKTAQDYKVQGGS--VLHLVLALRGGLQ-------------
T.castaneum(D6WL56)                 ---MLIKVKTLTGKEIEI-DIEPTDKVERIKERVEEKEGIPPQQQRLIFSGKQMNDEKTAQDYKVQGGS--VLHLVLALRGGTSF------------
P.humanus(E0VU21)                   ---MLIKVKTLTGKEIEI-DIEPTDKVERIKERVEEKEGIPPQQQRLIFSGKQMNDEKTAQDYKVQGGS--VLHLVLALRGGYSNIISAIN------
R.pedestris(R4WQJ5)                 ---MLIKVKTLTGKEIEI-DIEPTDKVERIKERVEEKEGIPPQQQRLIFSGKQMNDEKTAADYKVQGGS--VLHLVLALRGGHNVSH----------
P.albipes(T1DFZ4)                   ---MLIKVKTLTGKEIEI-DIEPTDKVDRIKERVEEKEGIPPQQQRLIFSGKQMNDDKTAQDYKVQGGS--VLHLVLALRGGHQ-------------
A.gambiae(Q7Q3J6)                   ---MLIKVKTLTGKEIEI-DIEPTDKVDRIKERVEEKEGIPPQQQRLIFSGKQMNDDKTAQDYKVQGGS--VLHLVLALRGGLLL------------
A.albopictus(A0A023EDB2)            ---MLIKVKTLTGKEIEI-DIEPTDKVDRIKERVEEKEGIPPQQQRLIFSGKQMNDDKTAQDYKVQGGS--VLHLVLALRGGRL-------------
D.melanogaster(X2J935)              ---MLIKVKTLTGKEIEI-DIEPTDKVDRIKERVEEKEGIPPQQQRLIFSGKQMNDDKTAADYKVQGGS--VLHLVLALRGGDSILTPCV-------
D.citri(A0A1S3DU76)                 ---MLIKVKTLTGKEIEI-DIEPTDKVERIKERVEEKEGIPPQQQRLIFSGKQMNDEKTAAEYKVQGGS--VLHLVLALRGGHSVIQSSLDQVLL--
R.appendiculatus(A0A131Z921)        ---MLIKVKTLTGKEIEI-DIEPNDKVERIKERVEEKEGIPPAQQRLIYSGKQMNDEKTATEYKVQGGS--VLHLVLALRGGQWR------------
L.unguis(A0A1S3KDM3)                ---MLIKVKTLTGKEIEI-DIEPTDKVERIKERVEEKEGIPPQQQRLIFSGKQMNDDKTAADYKVQGGS--VLHLVLALRGGHRGH-----------
S.pistillata(A0A2B4RW20)            ---MLIKVKTLTGKEIEI-DIEPTDKVERIKERVEEKEGIPPPQQRLIFSGKQMNDDKTAADYKIQGGS--VLHLVLALRGGLF-------------
C.horridus(A0A0K8RYZ5)              ---MLIKVKTLTGKEIEI-DIEPTDKVERIKERVEEKEGIPPQQQRLIYSGKQMNDEKTAADYKIQGGS--VLHLVLALRGGLR-------------
O.mykiss(C1BFL8)                    ---MLIKVKTLTGKEIEI-DIEPTDKVERIKERVEEKEGIPPQQQRLIYSGKQMNDEKTAADYKIQGGS--VLHLVLALRGGLVCHCNRIQLIV---
S.salar(B5X8K6)                     ---MLIKVKTLTGKEIEI-DIEPTDKVERIKERVEEKEGIPPQQQRLIYSGKQMNDEKTAADYKIQGGS--VLHLVLALRGGEVLHCPTSLLLAL--
G.aculeatus(G3PMC3)                 ---MLIKVKTLTGKEIEI-DIEPTDKVERIKERVEEKEGIPPQQQRLIYSGKQMNDEKTAADYKIQGGS--VLHLVLALRGGSRPRR----------
M.mola(A0A3Q4BER3)                  ---MLIKVKTLTGKEIEI-DIEPTDKVERIKERVEEKEGIPPQQQRLIYSGKQMNDEKTAADYKIQGGS--VLHLVLALRGGSTLRRPCTRLSPTS-
X.couchianus(A0A3B5LLA6)            ---MLIKVKTLTGKEIEI-DIEPTDKVERIKERVEEKEGIPPQQQRLIYSGKQ-NDEKTAADYKIQGGS--VLHLVLALRGGSEPH-----------
P.reticulata(A0A3P9QEH4)            ---MLIKVKTLTGKEIEI-DIEPTDKVERIKERVEEKEGIPPQQQRLIYSGKQMNDEKTAADYKIQGGS--VLHLVLALRGGSEPHRSCTRLSPSS-
P.mexicana(A0A3B3YIE0)              ---MLIKVKTLTGKEIEI-DIEPTDKVERIKERVEEKEGIPPQQQRLIYSGKQMNDEKTAADYKIQGGS--VLHLVLALRGGSEPHRSCTRLSPSS-
P.latipinna(A0A3B3UNR2)             ---MLIKVKTLTGKEIEI-DIEPTDKVERIKERVEEKEGIPPQQQRLIYSGKQMNDEKTAADYKIQGGS--VLHLVLALRGGSEPHRSCTRLSPSS-
T.nigroviridis(H3CR82)              ---MLIKVKTLTGKEIEI-DIEPTDKVERIKERVEEKEGIPPQQQRLIYSGKQMNDEKTAADYKIQGGS--VLHLVLALRGGSPS------------
A.limnaeus(A0A2I4AWV4               ---MLIKVKTLTGKEIEI-DIEPTDKVERIKERVEEKEGIPPQQQRLIYSGKQMNDEKTAADYKIQGGS--VLHLVLALRGGSPPRSCSTHLSSSS-
A.percula(A0A3P8TSK8)               ---MLIKVKTLTGKEIEI-DIEPTDKVERIKERVEEKEGIPPQQQRLIYSGKQMNDEKTAADYKIQGGS--VLHLVLALRGGSELHSRCIHVSSSS-
L.chalumnae(H3A9S3)                 ---TIAKVQTLTGKEIEI-DIEPTDKVERIKERVEEKEGIPPQQQRLIYSGKQMNDEKTAADYKIQGGS--VLHLVLALRGGSR-------------
E.kenyoni(A0A2Y9IJM8)               ---MLIKVKTLTGKEIEI-DIEPTDKVERIKERVEEKEGIPPQQQRLIYSGKQMNDEKTAADYKILGGS--VLHLVLALRGGGGLRQ----------
F.catus(M3W4C3)                     ---MLIKVKTLTGKEIEI-DIEPTDKVERIKERVEEKEGIPPQQQRLIYSGKQMNDEKTAADYKILGGS--VLHLVLALRGGGGLRQ----------
```

```
C.lupus(F1P890)                 ---MLIKVKTLTGKEIEI-DIEPTDKVERIKERVEEKEGIPPQQQRLIYSGKQMNDEKTAADYKILGGS--VLHLVLALRGGGGLRQ----------
E.europaeus(A0A1S3AKE6)         ---MLIKVKTLTGKEIEI-DIEPTDKVERIKERVEEKEGIPPQQQRLIYSGKQMNDEKTAADYKILGGS--VLHLVLALRGGGGLRQ----------
E.caballus(A0A3Q2HYF4)          ---MLIKVKTLTGKEIEI-DIEPTDKVERIKERVEEKEGIPPQQQRLIYSGKQMNDEKTAADYKILGGS--VLHLVLALRGGGGLRQ----------
C.porcellus(A0A286XT26)         ---MLIKVKTLTGKEIEI-DIEPTDKVERIKERVEEKEGIPPQQQRLIYSGKQMNDEKTAADYKILGGS--VLHLVLALRGGGGLRQ----------
P.coquereli(A0A2K6F9I7)         ---MLIKVKTLTGKEIEI-DIEPTDKVERIKERVEEKEGIPPQQQRLIYSGKQMNDEKTAADYKILGGS--VLHLVLALRGGGGLRQ----------
C.angolensis(A0A2K5IA72)        ---MLIKVKTLTGKEIEI-DIEPTDKVERIKERVEEKEGIPPQQQRLIYSGKQMNDEKTAADYKILGGS--VLHLVLALRGGGGLRQ----------
U.maritimus(A0A384DJ45)         ---MLIKVKTLTGKEIEI-DIEPTDKVERIKERVEEKEGIPPQQQRLIYSGKQMNDEKTAADYKILGGS--VLHLVLALRGGGGLRQ----------
P.macrocephalus(A0A2Y9FV62)     ---MLIKVKTLTGKEIEI-DIEPTDKVERIKERVEEKEGIPPQQQRLIYSGKQMNDEKTAADYKILGGS--VLHLVLALRGGGGLRQ----------
O.divergens(A0A2U3W261)         ---MLIKVKTLTGKEIEI-DIEPTDKVERIKERVEEKEGIPPQQQRLIYSGKQMNDEKTAADYKILGGS--VLHLVLALRGGGGLRQ----------
U.horribilis(A0A3Q7VHA4)        ---MLIKVKTLTGKEIEI-DIEPTDKVERIKERVEEKEGIPPQQQRLIYSGKQMNDEKTAADYKILGGS--VLHLVLALRGGGGLRQ----------
V.vulpes(A0A3Q7TQ17)            ---MLIKVKTLTGKEIEI-DIEPTDKVERIKERVEEKEGIPPQQQRLIYSGKQMNDEKTAADYKILGGS--VLHLVLALRGGGGLRQ----------
C.ursinus(A0A3Q7NXY2)           ---MLIKVKTLTGKEIEI-DIEPTDKVERIKERVEEKEGIPPQQQRLIYSGKQMNDEKTAADYKILGGS--VLHLVLALRGGGGLRQ----------
N.schauinslandi(A0A2Y9HNT6)     ---MLIKVKTLTGKEIEI-DIEPTDKVERIKERVEEKEGIPPQQQRLIYSGKQMNDEKTAADYKILGGS--VLHLVLALRGGGGLRQ----------
N.asiaeorientalis(A0A341C3R4)   ---MLIKVKTLTGKEIEI-DIEPTDKVERIKERVEEKEGIPPQQQRLIYSGKQMNDEKTAADYKILGGS--VLHLVLALRGGGGLRQ----------
H.sapiens(Q15843)               ---MLIKVKTLTGKEIEI-DIEPTDKVERIKERVEEKEGIPPQQQRLIYSGKQMNDEKTAADYKILGGS--VLHLVLALRGGGGLRQ----------
L.pardinus(A0A485MAU2)          ---MLIKVKTLTGKEIEI-DIEPTDKVERIKERVEEKEGIPPQQQRLIYSGKQMNDEKTAADYKILGGS--VLHLVLALRGGGGFRQ----------
N.vison(U6CT53)                 ---MLIKVKTLTGKEIEI-DIEPTDKVERIKERVEEKEGIPPQQQRLIYSGKQMNDEKTAADYKILGGS--VLHLVLALRGGGDLRQ----------
P.abelii(H2NKV7)                ---MLIKVKTLTGKEIEI-DIEPTDKVERIKERVEEKEGISPQQQRLIYSGKQMNDEKTAADYKILGGS--VLHLVLALRGGGGLRQ----------
M.auratus(A0A1U8CYD2)           ---MLIKVKTLTGKEIEI-DIEPTDKVERIKERVEEKEGIPPQQQRLIYSGKQMNDEKTAADYKILGGS--VLHLVLALRGGGGLGQ----------
C.griseus(G3HDD4)               ---MLIKVKTLTGKEIEI-DIEPTDKVERIKERVEEKEGIPPQQQRLIYSGKQMNDEKTAADYKILGGS--VLHLVLALRGGGGLGQ----------
I.tridecemlineatus(qI3N6M7)     ---MLIKVKTLTGKEIEI-DIEPTDKVERIKERVEEKEGIPPQQQRLIYSGKQMNDEKTAADYKILGGS--VLHLVLALRGGDSLRQ----------
G.gorilla(G3RJ38)               ---------TLTGKEIEI-DIEPTDKVERIKERVEEKEGIPPQQQRLIYSGKQMNDEKTAADYKILGGS--VLHLVLALRGGGGLRQ----------
A.melanoleuca(D2HG47)           --------QTLTGKEIEI-DIEPTDKVERIKERVEEKEGIPPQQQRLIYSGKQMNDEKTAADYKILGGS--VLHLVLALRGGGGLRQ----------
O.aries(W5QCY9)                 -------FQTLTGKEIEI-DIEPTDKVERIKERVEEKEGIPPQQQRLIYSGKQMNDEKTAADYKILGGS--VLHLVLALRGGGGLRQ----------
U.americanus(A0A452SHB5)        ---NPCLFQTLTGKEIEI-DIEPTDKVERIKERVEEKEGIPPQQQRLIYSGKQMNDEKTAADYKILGGS--VLHLVLALRGGGGLRQ----------
A.mississippiensis(A0A151M8R3)  ---MLIKVKTLTGKEIEI-DIEPTDKVERIKERVEEKEGIPPQQQRLIYSGKQMNDEKTAADYKIQGGS--VLHLVLALRGGGLR------------
G.gallus(A0A1D5PRI6)            ---MLIKVKTLTGKEIEI-DIEPTDKVERIKERVEEKEGIPPQQQRLIYSGKQMNDEKTAADYKIQGGS--VLHLVLALRGGGGVKQWGRF------
C.milii(V9LK01)                 ---MLIKVKTLTGKEIEI-DIEPTDKVERIKERVEEKEGIPPQQQRLIYSGKQMNDEKTASDYKIQGGS--VLHLVLALRGGAGL------------
M.domestica(F6T1M0)             ---MLIKVKTLTGKEIEI-DIEPTDKVERIKERVEEKEGIPPQQQRLIYSGKQMNDEKTAADYKIQGGS--VLHLVLALRGGGGLGR----------
R.norvegicus(A0A0G2K4G3)        ---MVLEQKTLTGKEIEI-DIEPTDKVERIKERVEEKEGIPPQQQRLIYSGKQMNDEKTAADYKILGGS--VLHLVLALRGGGGLGQ----------
L.africana(G3U8Y0)              ---MFIFQKTLTGKEIEI-DIEPTDKVERIKERVEEKEGIPPQQQRLIYSGKQMNDEKTAADYKILGGS--VLHLVLALRGGGGIRK----------
M.zebra(A0A3P9D203)             ---MLIKVKTLTGKEIEI-DIEPTDKVERIKERVEEKEGIPPQQQRLIYSGKQMNDEKTAADYKIQGGS--VLHLVLALRGGCLLHHRTTLCSTPAV
S.dumerili(A0A3B4TTJ4)          ---MLIKVKTLTGKEIEI-DIEPTDKVERIKERVEEKEGIPPQQQRLIYSGKQMNDEKTAADYKIQGGS--VLHLVLALRGGQVLHSPRCLYSKPAL
H.comes(A0A3Q2Y7I5)             ---MLIKVKTLTGKEIEI-DIEPTDKVERIKERVEEKEGIPPQQQRLIYSGKQMNDEKTAADYKIQGGS--VLHLVLALRGGKAHHTPDVQLASS--
D.rerio(F1QMF9)                 ---MLIKVKTLTGKEIEI-DIEPTDKVERIKERVEEKEGIPPQQQRLIYSGKQMNDEKTAADYKIQGGS--VLHLVLALRGGQIHQKPDRLLLNM--
A.mexicanus(A0A3B1JL70)         ---MLIKVKTLTGKEIEI-DIEPTDKVERIKERVEEKEGIPPQQQRLIYSGKQMNDEKTASDYKIQGGS--VLHLVLALRGGRIIY-----------
I.punctatus(E3TDT0)             ---MLIKVKTLTGKEIEI-DIEPTDKVERIKERVEEKEGIPPQQQRLIYSGKQMNDEKTAADYKIQGGS--VLHPVLALRGGRVHQHPNNLLQAL--
P.nattereri(A0A3B4EDC6)         ---MLIKVKTLTGKEIEI-DIEPTDKVERIKERVEEKEGIPPQQQRLIYSGKQMNDEKTAADYKIQGGS--VLHLVLALRGGREHQHPRNHLQAL--
E.lucius(C1BY35)                ---MLIKVKTLTGKEIEI-DIEPTDKVERIKERVEEKEGIPPQQQRLIYSGKQMNDEETAADYKIQGGS--VLHLVLAPRGGLLYHCPSIQIHS---
M.albus(A0A3Q3QM89)             ---MLETFATLTGKEIEI-DIEPTDKVERIKERVEEKEGIPPQQQRLIYSGKQMNDEKTAADYKIQGGS--VLHLVLALRGGSVLHGPCCIFFSLS-
L.loa(A0A1I7W0I3)               ---MLIKVKTLTGKEIEL-DIEPNDKVERIKEKVEEKEGIPPPQQRLIFAGKQMNDEKTAQDYKVAGGS--VLHLVLALRGGA--------------
A.grahami(A0A3N0XJQ5)           ---MHLFLKTLTGKEIEI-DIEPTDKVERIKERVEEKEGIPPQQQRLIYSGKQMNDEKTAADYKIQGGS--VLHLVLALRGGRLHQHLRSLLLTM--
L.corymbifera(A0A068SBG6)       ---MLIKVKTLTGKEIDI-DIEPTDKISRIKERVEEKEGIPPQQQRLIFSGKQMADEMTANEYSIVGGS--VLHLVLALRGGN--------------
R.microsporus(A0A0A1N8D5)       ---MLVKVKTLTGKEIEI-DIEPSDKISRIKERVEEKEGIPPQQQRLIYSGKQMSDEMTASQYGIEGGS--VLHLVLALRGGQ--------------
```

```
C.sinensis(H2KVL7)                 ---MLIKVKTLTGKEIEI-DIEPTDKVERIKERIEEKEGIPPPQQRLIFSGKQMHDEKVVSDYKIQGGS--VIHLVLSLRGGSR-------------
E.multilocularis(A0A068YBH6)       ---MLIKVKTLVGKEIEI-DIDQTDKIERIKERIEEKEGIPPPQQRLIFSGKQMNDEKTVNDYKIQGGS--VLHLVLALRGGQKH------------
D.discoideum(Q54XV3)               ---MLIKVKTLTGKEIEI-DIDPTDKIQRIKERVEEKEGIPPSQQRLIFGGKQMGDDKPASEYSIEGGS--VLHLVLALRGGL--------------
S.reilianum(E6ZVZ6)                ---MLIKVKTLTGKEIEL-DIEQTDKIQRIKERVEEKEGIPPAQQRLIFGGKQMHDEKTAKDFGVEGGS--VLHLVLALRGGSL-------------
A.ingoldii(A0A316YTF5)             ---MLIKVKTLTGKEIEL-DIEPTDKITRIKERVEEKEGIPPAQQRLIFGGKQMHDEKTAKEFGVEGGS--VLHLVLALRGGAFVGAMRG-------
A.amnicola(A0A1D1YNW4)             ---TMIKVKTLTGKEIEI-DIEPTDTIDRIKERVEEKEGIPPIQQRLIYAGKQLADDKTAKEYNIEGGS--VLHLVLALRGGLLS------------
M.truncatula(G7K8J5)               ---TMIKVKTLTGKEIEI-DIEPTDTIDRIKERVEEKEGIPPVQQRLIYAGKQLADDKTAKEYNIEGGS--VLHLVLALRGGN--------------
G.soja(A0A445GYG4)                 ---TMIKVKTLTGKEIEI-DIEPTDTIDRIKERVEEKEGIPPVQQRLIYAGKQLADDKTAKEYNIEGGS--VLHLVLALRGGTY-------------
V.radiata(A0A1S3U8Q0)              ---TMIKVKTLTGKEIEI-DIEPTDTIDRIKERVEEKEGIPPVQQRLIYAGKQLADDKTAKEYNIEGGS--VLHLVLALRGGIY-------------
D.antarctica(P0C073)               ---TMIKVKTLTGKEIEI-DIEPTDTIDRVKERVEEKEGIPPVQQRLIYAGKQLADDKTAKDYNIEGGS--VLHLVLALRGGY--------------
T.urartu(M7ZY92)                   ---TMIKVKTLTGKEIEI-DIEPTDTIDRIKERVEEKEGIPPVQQRLIYAGKQLADDKTAKDYNIEGGS--VLHLVLALRGGY--------------
P.dactylifera(A0A2H3X0H4)          ---TMIKVKTLTGKEIEI-DIEPTDTIDRIKERVEEKEGIPPVQQRLIYAGKQLADDKTAKDYNIEGGS--VLHLVLALRGGL--------------
B.rapa(V5M3G8)                     ---TMIKVKTLTGKEIEI-DIEPTDTIDRIKERVEEKEGIPPVQQRLIYAGKQLADDKTAKDYNIEGGS--VLHLVLALRGGFGLL-----------
S.lycopersicum(AOA3Q7I9N5)         ---TMIKVKTLTGKEIEI-DIEPTDTIDRIKERVEEKEGIPPVQQRLIYAGKQLADDKTAKDYNIEGGS--VLHLVLALRGGRL-------------
N.sylvestris(A0A1U7YZI3)           ---TMIKVKTLTGKEIEI-DIEPNDTIDRIKERVEEKEGIPPVQQRLIYAGKQLADDKTAKDYNIEGGS--VLHLVLALRGGRL-------------
S.splendens(A0A4D8ZGG0)            ---TMIKVKTLTGKEIEI-DIEPTDTIDRIKERVEEKEGIPPVQQRLIYAGKQLGDDKTAKDYNIEGGS--VLHLVLALRGGSH-------------
J.regia(A0A2I4E887)                ---TMIKVKTLTGKEIEI-DIEPTDTIDRIKERVEEKEGIPPVQQRLIYAGKQLGDDKTAKDYNIEGGS--VLHLVLALRGGSI-------------
C.melo(A0A1S3C469)                 ---TMIKVKTLTGKEIEI-DIEPTDTIDRIKERVEEKEGIPPVQQRLIYAGKQLGDDKTAKDYNIEGGS--VLHLVLALRGGHL-------------
G.arboreum(A0A0B0N165)             ---TMIKVKTLTGKEIEI-DIEPTDTIDRIKERVEEKEGIPPVQQRLIYAGKQLADDKTARDYNIEGGS--VLHLVLALRGGCL-------------
N.nucifera(A0A1U7ZNS2)             ---TMIKVKTLTGKEIEI-DIEPTDTIERIKERVEEKEGIPPVQQRLIYAGKQLADDKTARDYNIEGGS--VLHLVLALRGGSL-------------
Z.marina(A0A0K9P544)               ---TMIKVKTLTGKEIEI-DIEPNDTIERIKERVEEKEGIPPVQQRLIYAGKQLSDDKTAKDYNIEGGS--VLHLVLALRGGGN-------------
A.macrogynus(A0A0L0S2I6)           ---MQIKVKTLTGKEIEI-DVEIDDKLTRVKEKVEEKEGIPPAQQRLIFGGKQMSDEKTLADYNIQGGS--VLHLVLSLRGGHC-------------
B.mutus(L8HUC3)                    ---MLIKVKTLTRKEIEI-DTEPTDKVEQIKERVKEKEGIPPEQQRLTYSGRQINVKKTSADYKLLSGS--VLHLVLALRGGSDRPCSILPPCP---
B.prasinos(K8F2I6)                 ---MMIKVKTLTGKEVEL-DVEVSDTIVRIKERMEEKEGIPPIQQRLIFGGKQMHDEKTAKDYNIEGGS--VIHMVLALRGGGCV------------
C.lustricola(A0A2T3AIS2)           ---MQIKVRTLTGKEIEL-DIEGNYLISQIKEQVEEKEGIPPVQQRLIYQGKQMADEKKASEYDLQGGN--VLHLVLALRGGQ--------------
G.clavigera(F0XRI8)                ---MLIKVRTLTGKEIEL-DIDSDYKVSQIKEKVEEKEGIPPVQQRLIYGGKQMVDDKTASDYALEGGA--TLHLVLALRGGL--------------
M.bolleyi(A0A136IM41)              ---MLIKVRTLTGKEIEL-DIEADYKVSQIKEKVEEKEGIPPVQQRLIYGGKQMVDDKSAADYALEGGA--TLHLVLALRGGSQ-------------
C.orbiculare(N4V966)               ---MLIKVRTLTGKEIEL-DIENEYKVSQIKEKVEEKEGIPPVQQRLIYGGKQMVDDKSASEYALEAGA--TLHLVLALRGGCSPDGF---------
C.fructicola(L2G896)               ---MLIKVRTLTGKEIEL-DIENEYKVSQIKEKVEEKEGIPPVQQRLIYGGKQMVDDKSASEYGLEAGA--TLHLVLALRGGKASDVC---------
C.adophora(A0A2V1C054)             ---MLIKVRTLTGKEIEL-DIEPDYKVSQIKEKVEEKEGIPPVQQRLIFGGKQMADEKTATEYALEGGA--TLHLVLALRGGVCL------------
U.pustulata(A0A1W5CZ69)            ---MNIKVRTLTGKEIEL-DIEPDYKVARIKERVEEKEGIPPVQQRLIYGGKQMADDKTAAEYSLEGGA--TLHLVLALRGGR--------------
L.maculans(E4ZWL8)                 ---MQIKVRTLTGKEIEL-DIEADYKVSRIKERVEEKEGIPPAQQRLIYGGKQMADDKTAADYQLEGGA--TLHLVLALRGGC--------------
A.calidoustus(A0A0U5FUQ5)          ---MLIKVRTLTGKEIEL-DIEADYKVSRIKERVEEKEGIPPVQQRLIFGGKQMADDKTAQDYNLEGGA--TLHLVLALRGGCDAL-----------
G.moniliformis(W7LQF3)             ---MLIKVRTLTGKEIEL-DIESDYKVSQIKEKVEEKEGIPPVQQRLIHGGKQMTDDKTAAEYNLSAGD--TLHLVLALRGGRWMA-----------
F.oxysporum(W9II10)                ---MLIKVRTLTGKEIEL-DIESDYKVSQIKEKVEEKEGIPPVQQRLIHGGKQMTDDKTAAEYNLSAGD--TLHLVLALRGGRWMA-----------
T.guizhouense(A0A1T3CLM4)          ---MLIKVRTLTGKEIEL-DIESDYKVSQIKEKVEEKEGIPPVQQRLIHGGKQMTDDKTAAEYNLVAGD--TLHLVLALRGGRHFQ-----------
T.spiralis(A0A0V1BQN1)             MFKMLVKVKTLTGKEVEL-DIDASDLVERIKEKIEEKEGIPPAQQRLIHAGKQMSEGKTAAEYKITGGA--VLHLVLALRGGSL-------------
T.zimbabwensis(A0A0V1I5L9)         MLNMLVKVKTLTGKEVEL-DIDASDLVERIKEKIEEKEGIPPAQQRLIHAGKQMSEGKTAAEYKITGGA--VLHLVLALRGGSL-------------
T.richinella(A0A0V1BQN1)           ---MLVKVKTLTGKEVEL-DIDASDLVERIKEKIEEKEGIPPAQQRLIHAGKQMSEGKTAAEYKITGGAKTMAEFVLNLGYGSDDVL----------
A.thaliana(O65381)                 ---MEIKVKTLTEKQIDI-EIELTDTIERIKERIEEKEGIPPVHQRIVYTGKQLADDLTAKHYNLERGS--VLHLVLALRGGCC-------------
M.lychnidis(U5H805)                ---MNIKIKTLTGKEIDL-DIEPNDTVERIKVALEEKEGIPPAQQRLIFMGKAMPEEKTASDLKVTPGS--TLHLILSLRGGQC-------------
C.albicans(A0A1D8PJP6)             ---MQIKVKTLTGRDMPL-DVEPQDKIIRIKEMMEEKEGISPAQQRLIFNGSQLNDDQTVQEAGIQAGA--SLHLVLTLRGGN--------------
M.pulcherrima(A0A4P6XIP0)          ---MQVKVKTLTGRDIPV-DVEPTDKVIRIKEMMEEKEGIPPAQQRLIFNGSQLNDDATVEAAGIQPGT--SLHLVLTLRGGL--------------
R.necatrix(A0A1S7UJG8)             ---MMIKVRTLTGKLIEF-DVEPMDTIEDVKKRVEEKEGILPAQQRLIYSGKQMHDDVKLGEGGVVPDA--TLHLVLTLRGGASPRQW---------
```

```
S.cerevisiae(Q03919)            ---MIVKVKTLTGKEISV-ELKESDLVYHIKELLEEKEGIPPSQQRLIFQGKQIDDKLTVTDAHLVEGM--QLHLVLTLRGGN--------------
K.saulgeensis(A0A1X7R683)       ---MFIKVKTLTGKEIEV-ELQETDRIYHIKELLEQKEGIPPSQQRLIFQGKQINDEDSVGDAKLVDGM--QLHLVLTLRGGQ--------------
O.sativa(Q7XCL5)                MSMITVKVKTLTGKEVEV-SIEATETVARIKEQVEAAEGIPPPQQTLIYGGRQLADDMTAEMCDLRHGS--ELHLVLALRGGLL-------------
A.mphiamblys(A0A1J5WMY2)        ---MLIKVRTLTGAENEI-NIGADEKAEKIKRLVEDTDGIPMAQQRLLHNGRQVQDSKTLREIGVESGA--VLHLVIALRGG---------------
Nematocida(A0A177EH30)          ---MHIKIKTLEGTEVQL-EVDKAMKVKQVKELLQEKEAISIEQQRLIYGGKQLVDANTLETYGIQNND--VLHLVLALRGG---------------
P.knowlesi(A0A384LGS4)          ---MQILVKTLTGKRQSF-NFEPSSSVFQIKMAIEEKEGIDAKQIRLIYSGKQMHDDMKLSDYRVVPGS--TIHMILQLRGG---------------
P.vivax(A0A1G4HJQ8)             ---MQILVKTLTGKRQSF-NFEPSSSVFQIKMAIEEKEGIDAKQIRLIYSGKQMHDDMKLSDYRVVPGS--TIHMILQLRGG---------------
P.vivax_like(PVL_000121000)     ---MQILVKTLTGKRQSF-NFEPSSSVFQIKMAIEEKEGIDAKQIRLIYSGKQMHDDMKLSDYRVVPGS--TIHMILQLRGG---------------
P.inui(C922_00503)              ---MQILVKTLTGKRQSF-NFEPSSSVFQIKMAIEEKEGIDAKQIRLIYSGKQMHDDMKLSDYRVVPGS--TIHMILQLRGG---------------
P.fragile(AK88_00191)           ---MQILVKTLTGKRQSF-NFEPSSSVFQIKMAIEEKEGIDAKQIRLIYSGKQMHDDMKLSDYRVVPGS--TIHMILQLRGG---------------
P.cynomolgi(PCYB_142240)        ---MQILVKTLTGKRQSF-NFEPSSSVFQIKMAIEEKEGIDAKQIRLIYSGKQMHDDMKLSDYRVVPGS--TIHMILQLRGG---------------
P.gallinaceum(A0A1J1GYI6)       ---MQILVKTLTGKRQSF-NFEPSSSVYQIKMAIEEKEGIDAKQIRLIYSGKQMHDDMKLSDYRVVPGS--TIHMILQLRGGII-------------
P.relictum(A0A1J1HAD2)          ---MQILVKTLTGKRQSF-NFEPSSSVYQIKMAIEEKEGIDAKQIRLIYSGKQMHDDMKLSDYRVVPGS--TIHMILQLRGGII-------------
P.falciparum(Q8IEI4)            ---MQILVKTLTGKRQSF-NFEPSSSVFQIKMAIEEREGIDAKQIRLIYSGKQMHDDMKLSDYRVVPGS--TVHMILQLRGG---------------
P.billcollinsi(PBILCG01_1313400) ---MQILVKTLTGKRQSF-NFEPSSSVFQIKMAIEEREGIDAKQIRLIYSGKQMHDDMKLSDYRVVPGS--TVHMILQLRGG---------------
P.blacklocki(PBLACG01_1311100)  ---MQILVKTLTGKRQSF-NFEPSSSVFQIKMAIEEREGIDAKQIRLIYSGKQMHDDMKLSDYRVVPGS--TVHMILQLRGG---------------
P.reichenowi(PRCDC_1312000)     ---MQILVKTLTGKRQSF-NFEPSSSVFQIKMAIEEREGIDAKQIRLIYSGKQMHDDMKLSDYRVVPGS--TVHMILQLRGG---------------
P.praefalciparum(PPRFG01_1314900) ---MQILVKTLTGKRQSF-NFEPSSSVFQIKMAIEEREGIDAKQIRLIYSGKQMHDDMKLSDYRVVPGS--TVHMILQLRGG---------------
P.adleri(PADL01_1311900)        ---MQILVKTLTGKRQSF-NFEPSSSVFQIKMAIEEREGIDAKQIRLIYSGKQMHDDMKLSDYRVVPGS--TVHMILQLRGG---------------
P.gaboni(PGABG01_1311100)       ---MQILVKTLTGKRQSF-NFEPSSSVLQIKMAIEEREGIDAKQIRLIYSGKQMHDDMKLSDYRVVPGS--TVHMILQLRGG---------------
P.ovale(A0A1C3L4R7)             ---MQILVKTLTGKRQSF-NFEPSSSVYQIKMAIEEREGIDAKQIRLIYSGKQMHDEMKLSDYRVVPGS--TIHMILQLRGGIF-------------
P.yoelii(A0A077YAC8)            ---MQILVKTLTGKRQSF-NFEPSSSVLQIKMAIEEREGIDAKQIRLIFSGKQMHDDMKLSDYRVIPGS--TIHMILQLRGGTI-------------
P.vinckei(YYE_01216)            ---MQILVKTLTGKRQSF-NFEPSSSVLQIKMAIEEREGIDAKQIRLIFSGKQMHDDMKLSDYRVIPGS--TIHMILQLRGGRI-------------
P.chabaudi(A0A077TQI6)          ---MQILVKTLTGKRQSF-NFEPSSSVLQIKMAIEEREGIDAKQIRLIFSGKQMHDDMKLSDYRVIPGS--TIHMILQLRGGRI-------------
P.berghei(A0A509AZB5)           ---MQILVKTLTGKRQSF-NFEPSSSVLQIKMAIEEREGIDAKQIRLIFSGKQMHDDMKLSDYRVIPGS--TIHMILQLRGGTN-------------
P.malariae(A0A1C3L273)          ---MQILVKTLTGKRQSF-NFEPSSTVYQIKMAIEEREGIDAKQIRLIYSGKQMHDEMKLSDYKVVPGS--TIHMILQLRGGKY-------------
T.gondii(A0A125YX02)            ---MQILIKTLTGRKQSF-NFEPDNTVLHVKQALQEKEGIDVKQIRLIYSGKQMSDELKLSDYKVVPGC--TIHMVLQLRGGC--------------
H.hammondi(HHA_254600)          ---MQILIKTLTGRKQSF-NFEPDNTVLHVKQALQEKEGIDVKQIRLIYSGKQMSDELKLSDYKVVPGC--TIHMVLQLRGGC--------------
N.caninum(F0V9J4)               ---MQILIKTLTGRKQSF-NFEPDNTVLHVKQALQEKEGIDVKQIRLIYSGKQMSDELKLSDYKVVPGC--TIHMVLQLRGGC--------------
C.parvum(A3FPR3)                ---MQILVKTLTGKKQNF-NFEPENTVLQVKQALQEKEGIDVKQIRLIYSGKQMSDDLRLLDYKVTAGC--TIHMVLQLRGGLR-------------
C.cayetanensis(A0A1D3D9Z9)      ---MQILVKTLTGRRQSF-NFEGENLVVHVKQALQEKEGVDVEQIRLIFSGKQMNDELKLSDYRVSPGC--TIHMVLQLRGGQSH------------
E.histolytica(C4LVJ5)           ---MLINIKLLNGRILSI-DLEPTDKISDLKAKLEEIEGITPEQQRLVFGGRQLGDDKTLQELNIQPGT--QINLLLALRGGF--------------
T.brucei(Q584D3)                ---MLLKVKTVSNKVIQITSLTDDNTIAELKGKLEESEGIPGNMIRLVYQGKQLEDEKRLKDYQMSAGA--TFHMVVALRAGC--------------
Consensus                          M-IKVKTLTGKE----D-EP---V--IKE--EEKEGIPP-QQRLI--GKQM-D-KT--DY----GS---LHLVLALRGG
                                                              **                                          *    ↑
```

Fig. S1

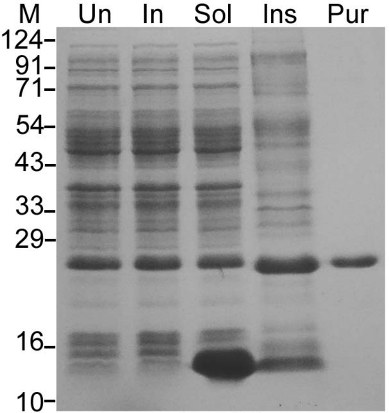


Fig. S2

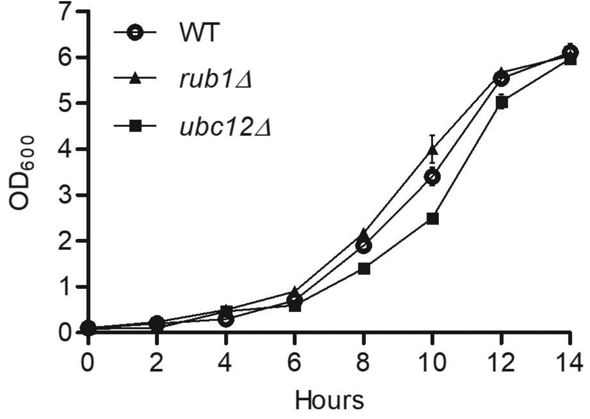


**Fig. S3**
